# The inhibitory control of traveling waves in cortical networks

**DOI:** 10.1101/2022.11.02.514832

**Authors:** Grishma Palkar, Jian-young Wu, Bard Ermentrout

## Abstract

Propagating waves of activity can be evoked and can occur spontaneously *in vivo* and *in vitro*. We examine the properties of these waves as inhibition varies in a cortical slice and then develop several computational models. We first show that in the slice, inhibition controls the velocity of propagation as well as the magnitude of the local field potential. We introduce a spiking model of sparsely connected excitatory and inhibitory theta neurons which are distributed on a one-dimensional domain and illustrate both evoked and spontaneous waves. The excitatory neurons have an additional spike-frequency adaptation current which limits their maximal activity. We show that increased inhibition slows the waves down and limits the participation of excitatory cells in this spiking network. Decreased inhibition leads to large amplitude faster moving waves similar to those seen in seizures. To gain further insight into the mechanism that control the waves, we then systematically reduce the model to a Wilson-Cowan type network using a mean-field approach. We simulate this network directly and by using numerical continuation to follow traveling waves in a moving coordinate system as we vary the strength and spread of inhibition and the strength of adaptation. We find several types of instability (bifurcations) that lead to the loss of waves and subsequent pattern formation. We approximate the smooth nonlinearity by a step function and obtain expressions for the velocity, wave-width, and stability.

**Author summary:** Stimuli and other aspects of neuronal activity are carried across areas in the brain through the concerted activity of recurrently connected neurons. The activity is controlled through negative feedback from both inhibitory neurons and intrinsic currents in the excitatory neurons. Evoked activity often appears in the form of a traveling pulse of activity. In this paper we study the speed, magnitude, and other properteis of these waves as various aspects of the negative feedback are altered. Inhibition enables information to be readily transmitted across distances without the neural activity blowing up into a seizure-like state.

## Introduction

Recordings in different cortical regions and layers during sensory stimulation show that the response often manifests as a traveling wave [1–5] even though the stimulus is localized. In these waves, groups of neurons fire transiently in succession and then falling to a depressed (refractory) state before returning to rest. Evoked waves are distinct from the waves that are seen in local field potentials of humans and animals which appear as spatio-temporal phase gradients in ongoing activity [1, 6]. They are more akin to traveling action-potentials in excitable media while the latter are more like the waves that arise from coupled oscillators [7]. For the evoked waves, the firing of excitatory cells is sparse (a characteristic of so-called balanced networks of excitation and inhibition [8]), yet the wave manages to propagate over centimeters of tissue. In contrast, when inhibition is pharmacologically blocked, particularly in slice preparations, the waves involve many excitatory neurons, have much broader profile (when imaged with voltage sensitive dyes) and travel faster [5, 9]. These reduced-inhibition waves have been implicated in seizure propagation models [10–12]. This suggests that the sparsity and degree of participation of excitatory cells in the propagation of evoked waves is controlled by negative feedback. (See also [13].) There are at least two sources to this negative feedback: recurrent inhibition and activity-dependent adaptation in the excitatory cells. Thus, our goal in this paper is to explore how these two effects work to control the macroscopic properties of evoked waves.

The earliest computational models for traveling waves (TWs) in neural media can be found in the the work of Wilson and Cowan [14] and Amari [15] who looked at dynamics in neural fields. Later models of TWs using Hodgkin-Huxley type spiking models with distance dependent coupling explored waves in thalamus [16] and disinhibited cortex [17]. [18, 19] provided some early attempts for the analysis of TWs in spiking models. Most of the analytic work on TWs has been done for neural field or firing rate models, where the activity of populations of neurons is modeled rather than individual cells. [10] analyzed a model which had a population of excitatory cells and a linear adaptation variable using a combination of analytic methods and singular perturbation theory. There have been many extensions of this work with comprehensive reviews found in [20, 21].

All of the above mentioned computational papers focused on either recurrent inhibition, spike frequency adaptation, or synaptic depression as means of preventing run away inflation. Only recently have models been suggested which involve multiple types of negative feedback. [12] analyze a system of excitatory and inhibitory rate models where the excitatory population includes adaptation. In their model, propagation occurs only when the inhibition is weak and slow. In the discussion, we will examine in more detail differences between their model and ours.

In this paper, we are interested in studying the effects of recurrent inhibition and spike frequency adaptation on the properties of evoked waves. We first show some stimulus evoked waves from rat cortical slices with intact inhibition and then when inhibition is blocked. Our first model is a network of spatially distributed sparsely coupled spiking excitatory and inhibitory theta neurons [22]. We vary the recurrent inhibition and study its effects on speed, wave failure, and fraction of neurons spiking. To get better insight, we next create an approximate mean field model by using a recently described connection between the theta neuron and exact mean field reductions [23, 24] and its generalization to space [25]. From this we are able to derive a Wilson-Cowan type neural field model. We use numerical continuation and simulations to study the effects of recurrent inhibition and adaptation on the velocity of evoked TWs. We find some interesting instabilities of the waves as the footprint of the inhibition increases allowing us to connect the failure to propagate with transition to stationary spatial patterns such as bump attractors and Turing patterns. Finally, we replace the smooth nonlinearities of the neural field model with step functions which enables us to obtain analytic results on the properties of the waves and their stability.

## Materials and methods

### Experimental methods

#### Cortical slice preparation

Sprague-Dawley rats of both sexes (P14-P35) and C57BL6 mice (P21-P60) were used in accordance with a protocol that had been approved by the Institutional Animal Care and Use Committee at Georgetown University Medical Center. Following deep isoflurane anesthesia, animals were rapidly decapitated. The whole brain was subsequently removed and chilled in cold (0°C) sucrose-based cutting artificial cerebrospinal fluid (sACSF) containing (in mM) 252 sucrose; 3 KCl; 2 CaCl_2_; 2 MgSO_4_; 1.25 NaH_2_PO_4_; 26 NaHCO_3_; 10 dextrose and bubbled by 95% O_2_, 5% CO_2_. Cortical slices (300 um thick) were cut in coronal sections from dorsal to ventral brain with a vibratome (Leica, VT1000S). Slices are incubated in ACSF contained (in mM) NaCl, 132; KCl, 3; CaCl_2_, 2; MgSO_4_, 2; NaH_2_PO_4_, 1.25; NaHCO_3_, 26; dextrose 10; and saturated with 95% O_2_, 5% CO_2_ at 26-27°C. The incubation chamber is equipped with a small pump to circulate oxygenated ACSF for 2-5 hours [26]. In well circulated conditions the slices can recover well from the injury of slicing. Fully recovered slices can remain viable for up to 24 hours [26, 27].

##### Field potential recording (LFP) in slice

Low resistance glass microelectrodes (50-150KΩ tip resistance) were used for LFP recordings. The electrodes were pulled with a Sutter P87 puller (Sutter Instruments) with 6 controlled pulls. Electrodes were filled with ACSF. The recordings were done in a submerged chamber, and slices were perfused on both sides at a high flow rate (10-30 ml/min). The LFP data are amplified 1000x by a custom-made amplifier (0.01-1000Hz) and digitized at 3000 Hz by a 12-bit USB Analog-to-digital converter (National instruments). From each brain slice we can usually record continuously for 3 - 8 hours.

##### Voltage-sensitive dye recording

The imaging apparatus and methods are described in detail in [28, 29]. Briefly, the slices were stained with ACSF containing 0.005 to 0.02 mg/ml of an oxonol dye, NK3630 [30] for 30 to 60 minutes. A 124-element photodiode array system was used for the imaging. The preparation was trans-illuminated by 705 ± 20 nm light for the imaging. An objective of 5 × (0.12 NA, Zeiss) was used to form the image on the diode array. Each photodetector received light from an area of 330 × 330 *μ*m*m*^2^ of the cortical tissue. With trans-illumination, neurons through the entire thickness of the slice(450 *μ*m) contributed equally to the signal. The resting light intensity was about 109 photons/msec per detector and the VSD signal of the oscillation was about 0.01% (peak-to-peak) the resting light intensity. The signal was AC coupled at 0.1 Hz, amplified 200 times, low-pass filtered at 333 Hz, and then digitized at 1,000 frames /sec with a 12 bit accuracy.

### Spiking Model

The spiking model consists of a network of *N_e_* = 400 excitatory (E) and *N_i_* = 80 inhibitory (I) quadratic integrate-and-fire neurons which we transform to a network of theta neurons for ease in integration. For simplicity of later reduction, we use “current” synapses rather than conductance-based synapses. Each excitatory neuron, *j* = 0,…, *N_e_* − 1 is positioned at a spatial location *x* ∈ [0, 1) where *x* = *j*/400 and each inhibitory neurons, *j* = 0,…, *N_i_* − 1 is positioned at *x* = *j*/80. Connections between neurons are made probabilistically and fixed:

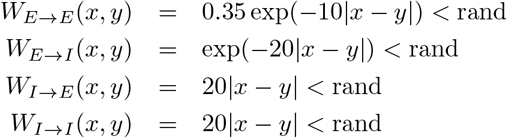

where rand is a uniform random number in (0, 1). This produces a fairly sparse distance-dependent network. Each E cell receives about 27 excitatory and 8 inhibitory inputs; each I cell receives about 40 excitatory and 8 inhibitory inputs. All inputs have weight 1. The voltage of each neuron evolves as:

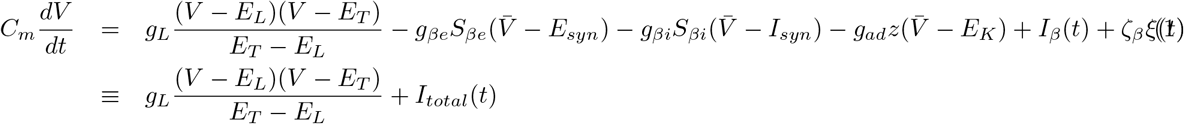

where *β* ∈ {*e, i*} and 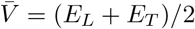. *S_βe_, S_βi_* are the weighted synaptic inputs. For example 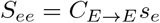. *z* is spike frequency adaptation and is only applied to the excitatory cells, with conducatnce, *g_ad_*. A neuron spikes at time *t_sp_* if lim 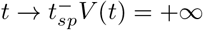 whence it is immediately reset to −∞. The synaptic variables, *s_e_, s_i_* and the adaptation, *z* satisfy first order kinetics:

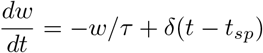

where *β* ∈ {*e, i, z*}. *ξ*(*t*) is Gaussian noise. Standard parameters are *C_m_* = 1*μF/cm*^2^, *g_L_* = 0.1*mS/cm*^2^, *g_ee_* = 0.15*mS/cm*^2^, *g_ei_* = 1*mS/cm*^2^, *g_ie_* = 0.4*mS/cm*^2^, *g_ii_* = 0.1*mS/cm*^2^, *E_L_*= −65*mV*, *E_T_*= −50*mV*, *E_K_* = −85*mV*, *E_syn_* = 0*mV*, *I_syn_* = −75*mV*, 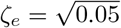, 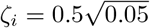, *τ_e_* = 3*msec*, *τ_i_* = 4*msec*, and *τ_z_* = 50*msec*. In order to simulate this model exactly, we transform it to the theta model:

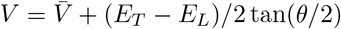

so that *θ* evolves as

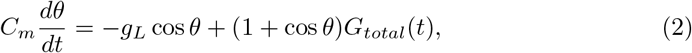

where *G_total_*(*t*) = 2*I_total_*(*t*)/(*E_T_* − *E_L_*) is the input conductance. Spiking occurs when *θ* crosses *π* from below; *θ* is then reset to −*π* and the corresponding synaptic and adaptation variables are incremented by 1. We integrate the system of equations with Euler-Maruyama method using dt = 0.05*msec*. We simulate the local field potential as:

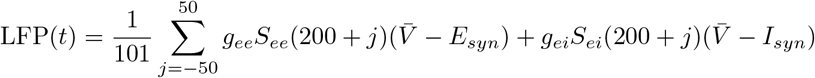

#### Reduction to firing rate model

In the results, we replace the Gaussian noise with a specific kind of heterogeneity, apply the methods of [24] and then assume a pseudo-steady state for the mean firing rate and potential to derive a firing rate function:

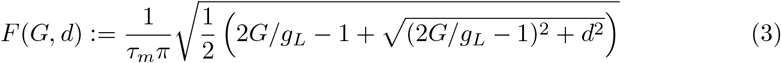

where *τ_m_* = 2*C_m_/g_L_* is twice the membrane time constant. The arguments to *f* are *G* = *G_total_* and *d* which proportional to the “noise” magnitude. We replace the random distance-dependent connectivity matrices with convolutions in space with some kernel (generally exponential, but also Gaussian). We thus consider the following set of “Wilson-Cowan”-like equations:

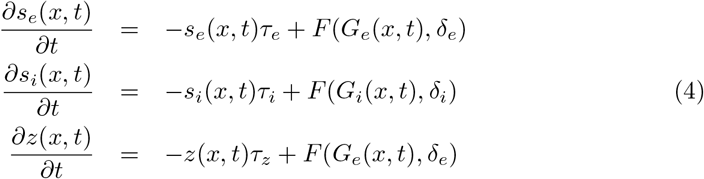

where

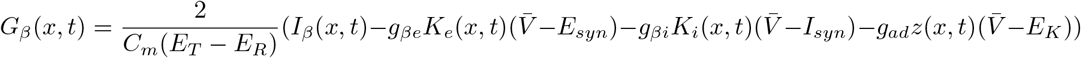

and *K_β_*(*x, t*) = *W_β_*(*x*) ⋆ *s_β_*(*x, t*) is the convolution of some kernel, *W* (*x*), with the synaptic activation, *s*(*x, t*). To solve this equation, we discretize the interval [0, 1] into 400 bins and solve the resulting set of ODEs using the Runge-Kutta algorithm with step size 0.05.

#### Traveling waves

Traveling waves satisfy, *s*(*x, t*) = *S*(*x* + *ct*) where *c* is the velocity of the wave and *s* ∈ {*s_e_, s_i_, z*}. If the convolution kernels, *W_e,i_*(*x*) are exponential, then they can be inverted to become second order ODEs in *x*. This means that the system Eq. (4) can be written as a 7-dimensional ODE (see results for details) with a 2-dimensional stable and 5-dimensional unstable manifold. We use XPPAUT to compute the homoclinic orbits for this system via continuation with AUTO. To get a starting approximation for AUTO, we first consider the case where *σ_i_* = 0 so that we have a 5-dimensional system. The equilibrium turns out to have a 1-dimensional unstable manifold which we approximate from the eigenvector of the positive eigenvalue. We can then vary *c* until we get close to a homoclinic orbit. We use this as a starting point and continue in *σ_i_* until we get to the default values of all the parameters.

#### Step function approximation

We approximate the smooth nonlinearity *F* with a scaled Heaviside step function and use this to derive a series of nonlinear equations for the properties of the waves such as the velocity, onset of inhibition and the width of the excitatory and inhibitory pulses. We use XPPAUT to study how these change with parameters. We formally linearize about the traveling waves and compute the so-called Evans Function [31] (see results for details). We then use MATLAB to find the roots of the Evans Function in order to assess the stability of the waves. We use the fsolve function in MATLAB to find the curves of Hopf bifurcations.

## Results

### Experimental results

To illustrate the effects of recurrent inhibition, we recorded evoked activity for a cortical slice preparation (see Materials & Methods for details). An electrical stimulation of moderate intensity (1-10V, 0.05 ms pulse) induced a local pulse of population activity propagating slowly though the cortex (Fig. 1A, right top trace). After removing inhibition with 20 *μ*M of bicuculline, the same stimulation induced a large and fast pulse of activity (Fig. 1A, bottom trace). Note that removing inhibition also largely reduced the latency of the evoked response. The latency included the time for the activity spreading from stimulation site to the recording site, and so the reduced latency suggested a faster propagating speed after removal of inhibition. The propagation velocity was further examined with multiple recording sites (Fig. 1B). In this experiment we used voltage-sensitive dye to convert membrane potentials to a light intensity signal. Each optical detector receives light from thousands of cortical neurons, and the optical signal is thus an integration of membrane potential change of all the neurons under one detector. We found that removing inhibition greatly increased the propagating speed of the cortical activity (> 20×). Our results have been verified by more than 100 slices since [32]. This example clearly shows several properties of the wave in normal and disinhibited slices. First, the response is substantially greater in both magnitude and duration when inhibition is removed. Further the wave shape is more reliable during the propagation event and far less noisy. Finally, the velocity changes dramatically. Our goal in the rest of this paper to to try to explain these properties via a computational model.

**Fig 1.**
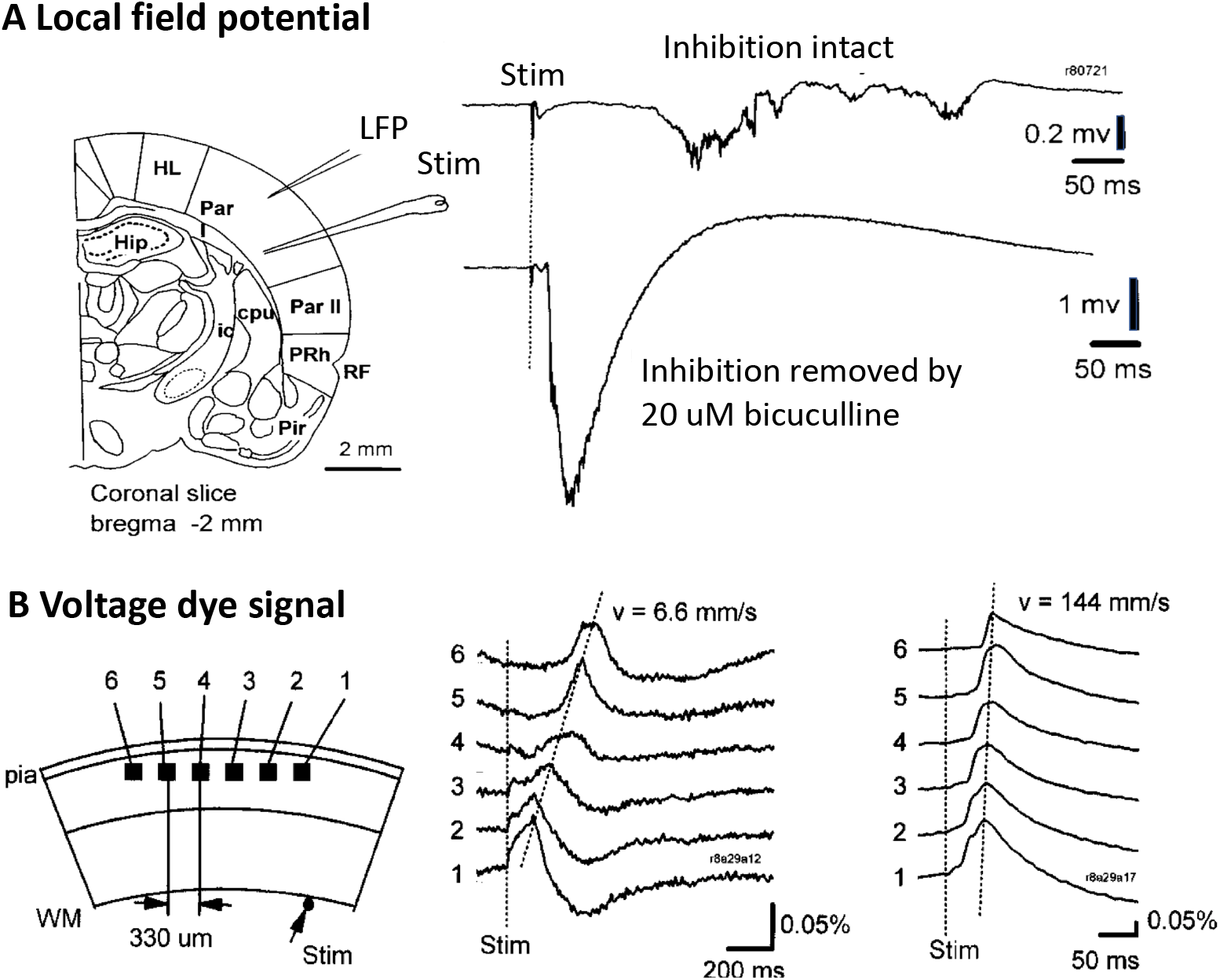
Evoked cortical activity and propagation velocity in a rat cortical slice. A. Neuronal activity measured by local field potential electrode. B. Propagating velocity measured by voltage-sensitive dye and optical recording. (Modified from [32])

### Spiking Model

We are interested in how recurrent inhibition controls aspects of waves that are generated by brief localized stimuli. In order to study this, we have created a network of 400 excitatory and 80 inhibitory neurons distributed on a one-dimnsional line 0 < *x* < 1 with random sparse distance-dependent connections (see Methods for details). Fig. 2 shows plots of *s_e_*(*j*), *s_i_*(*j*) for the network of 480 cells. At *t* = 20, a brief pulse of current is applied to the first 20 excitatory cells and evokes a wave. Only about 10% of the excitatory cells participate in the wave, where as nearly all the inhibitory cells do. (For this reason, it is easier to visually follow the wave by looking at the inhibitory response.) The wave takes about 50 msec to traverse the domain. As the recurrent inhibition (*g_ei_*) decreases, the number of excitatory cells participating in the wave increases dramatically, with nearly 100 % participation once *g_ei_* falls below 0.25. There is a small increase in velocity, but it is not as dramatic as shown in Fig. 1B. Increasing *g_ei_* beyond 1 does little to affect the speed or participation. Fig. 3A shows how the inhibition reduces participation of the excitatory cells in the wave. We show *θ*(*t*) (the transformed voltage) for a selected excitatory cell (#195) near the middle of the domain. Strong excitation from cell #184 produces a sufficient response such that without subsequent inhibition, the E cell would reach *θ* = *π* and spike. However, the inhibition from cell # 40 (also, in the middle of the domain) shuts this response down. Because of the large number of excitatory to inhibitory synapses in our model, the inhibitory cells have a high probability of firing and thus keep the fraction of excitatory cells that participate to a low number as suggested in [13, 33] Fig. 3B shows the fraction of excitatory cells participating in the wave as the parameter *g_ei_* changes. Because the connections are randomly drawn from a distribution and fixed throughout, the degree of participation has a bit of “noise”, but the overall trend is quite clear.

**Fig 2.**
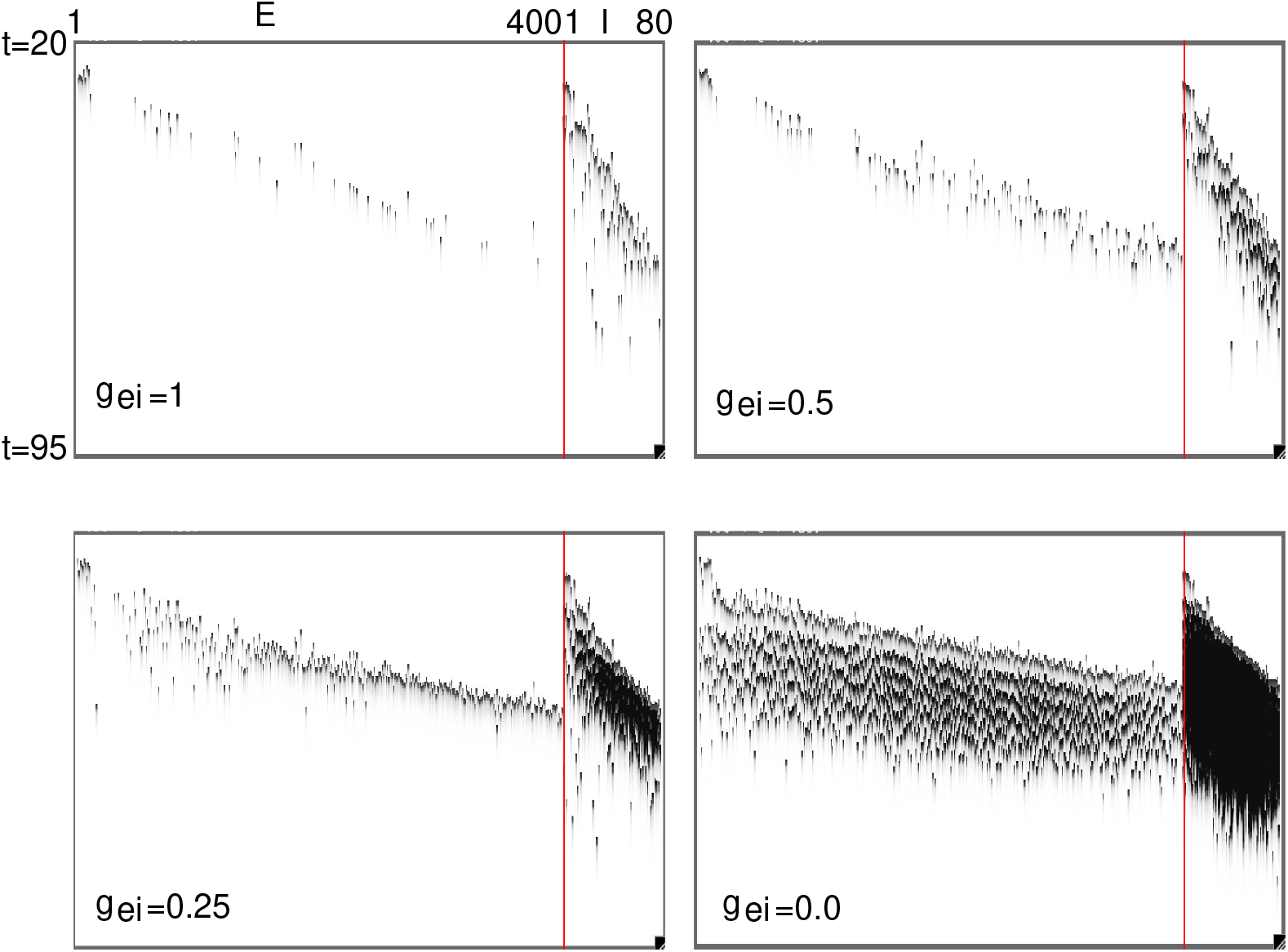
Traveling waves generated in the spiking models are controlled through the strength of feedback inhibition to the excitatory cells, *g_ei_*.

**Fig 3.**
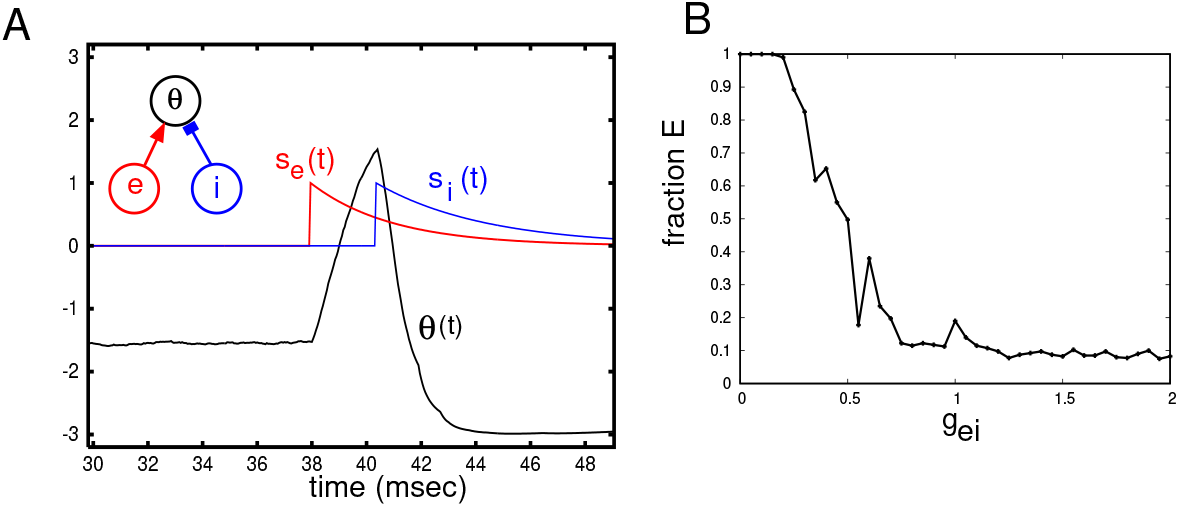
(A) Sparse coupling and feedforward inhibition control the participation in waves. *θ*(*t*) for excitatory cell #195 is excited by cell # 184, but before firing is inhibited by cell # 40. (B) Fraction of excitatory cells participating in the wave varies with *g_ei_*

In preparation for the ensuing mean field analysis, we plot some averaged quantities in Fig. 4. We average the excitatory and inhibitory synapses, *s_e_*(*j, t*), *s_i_*(*j, t*) for *j* in the middle of the network (*j* = 200 for the excitatory and *j* = 40 for the inhibitory cells). Panel A shows the evolution in time of 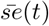 for four different values of *g_ei_*. Panel B shows the projection in the 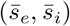 phase plane. As long as *g_ei_* is not too small, the mean inhibition is only 2-3 times larger than the mean excitation, but when *g_ei_* = 0, the very large excitation makes the inhibitory cells fire many spikes during the wave. Looking again at the raster plots, Fig. 2, one can see that even with high values of *g_ei_*, some inhibitory nwurons fire multiple times during the wave. The model LFP in Fig. 4C is qualitatively similar to the experimental trace in Fig. 1A. (We note that the model LFP is not a true LFP, but rather a sum of the excitatory and inhibitory currents into a group of model cells.)

**Fig 4.**
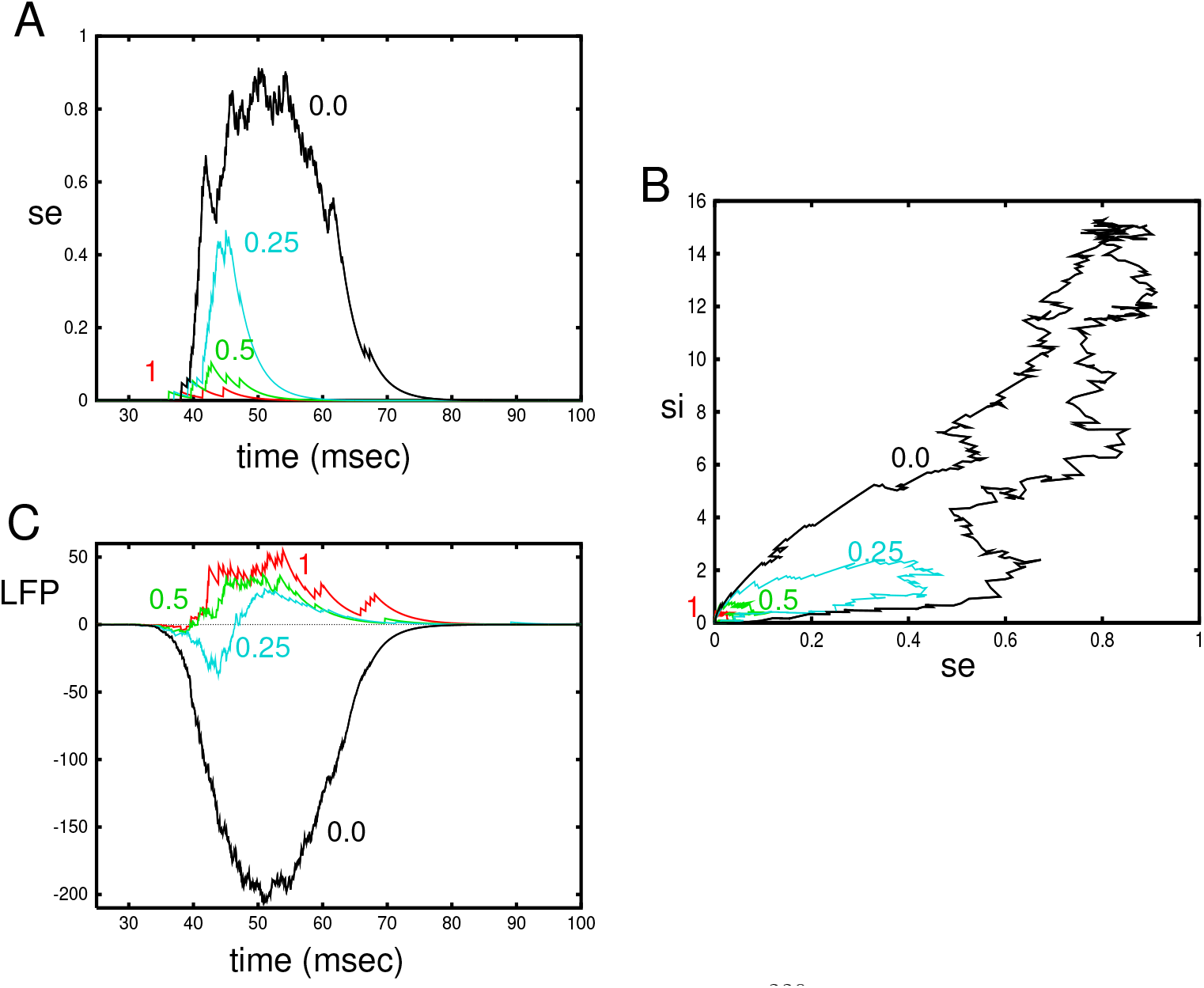
Averaged activity as *g_ei_* varies. (A) 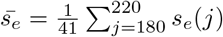 for 4 values of *g_ei_*. (B) Projections of 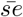 and 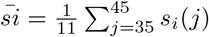 for different values of *g_ei_*. (C) “LFP” (see Methods) for different values of *g_ei_*

#### 0.1 Mean field reduction

The advantage of the *θ* model and the corresponding QIF model is that there is a rigorous mean field reduction (MFR) under certain circumstances [24]. Laing and others have extended these ideas to spatial networks in certain limits [25]. As we are ultimately not going to use the full MFR, but an even more reduced version, we only briefly sketch the derivation. We consider a local network (no space) of excitatory and inhibitory neurons with all-to-all coupling. Let *u* = tan(*θ*/2) be a dimensionaless voltage. Then for each population, Eq. (2) can be written as:

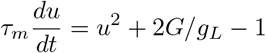

where *τ_m_* = 2*C_m_/g_L_* is twice the membrane time constant. (The factor of 2 comes from our original transformation of *V* to *θ*.) If we replace the Gaussian noise, *ξ_j_* in Eq. (1) by a random applied current taken from a Cauchy distribution 1/(*π*(1 + *ξ*^2^)), then, following [24] and others, we derive an equation for two quantities, (*η_β_, ν_β_*) (*β* ∈ {*e, i*}) that satisfy:

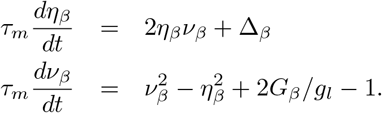

Here Δ_*β*_ = 2*ζ_β_*/(*g_L_*(*E_T_ − E_R_*)) where *ζ_β_* is the heterogeneity. The adaptation and synaptic inputs are replaced by their means and evolve as:

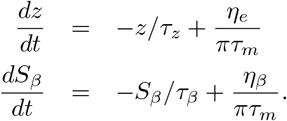

*η_β_*/(*πτ_m_*) is the mean firing rate of the population and 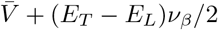 is the potential of the population. Finally,

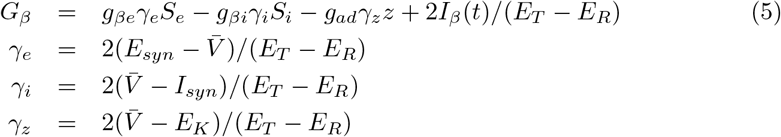

(Note: (i) there is no adaptation for the inhibitory population; (ii) we have rearranged *γ_e_* so that it has a positive sign as do *γ_i,z_*) If we want to extend this system to space, then each of the *η, ν* will also be space dependent as will the mean synapses and adaptation. This system of equations is an exact reduction of the full spiking model in the limit as the number of neurons goes to infinity and when the Gaussian noise is replaced by the Cauchy-distributed inputs. We could thus simulate this system of equations instead of the spiking model. But, the quadratic nonlinearities in the (*η, ν*) variables make integration difficult and the numerical shooting that we will eventually do very challenging. Thus, we make one final reduction and eliminate (*η, ν*) altogether. If we hold *G* constant (pseudo-steady state assumption), we can solve for the equilibrium solution for (*η, ν*): *ν* = −Δ/(2*η*) and *η* satisfies a quartic equation of the form *c*_2_*η*^4^ + *c*_1_*η*^2^ + *c*_0_ = 0 which we can solve using the quadratic formula applied to *η*^2^. The evolution of the mean synaptic variables and adaptation depend only on the firing rate, *η*/(*πτ_m_*) which we obtain from solving the quartic equation and thus recover Eq. (3).

### Local kinetics

With this reduction, we can now consider the “Wilson-Cowan” type model Eq. (4). The spiking simulations suggest a pulse of activity propagates across the medium. The hallmark of pulse waves is an underlying excitable medium. Thus, we focus on the excitability of the local (*s_e_*, *s_i_*, *z*) system. With two exceptions, we will use the same parameters for the mean field model as we used in the spiking model. In the spiking model, we chose *g_ee_* = 0.15, but for the mean field models we increase this to *g_ee_* = 1. We also reduce *g_ad_* = 1 to *g_ad_* = 0.25, although we will vary this parameter later in the paper. We consider

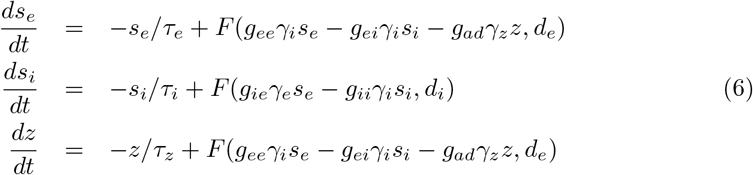

Figure 5A depicts the behavior for this three-dimensional system when *s_e_* is perturbed away from 0. The recurrent excitation causes an initial amplification of *s_e_*(*t*) before the inhibition and adaptation suppress it. We can better explore the nature of the excitability by setting either of *g_ei_* or *g_ad_* to zero leading to a planar system. In panel B (*g_ad_* = 0), we see that the nullclines intersect in three places. There is a single stable equilibrium near (*s_e_*, *s_i_*) = (0, 0), a saddle point near (0, 0.005) and an unstable node (blue circle). We superimpose a trajectory starting at (0, 0.01) which decays back to the rest state after an excursion in the plane. Thus, the (*s_e_, s_i_*) subsystem is characterized by Class I excitability [34]. In panel C, we allow the adaptation (*g_ad_* = 0.25) but set *g_ei_* = 0 and depict a typical trajectory starting at *z* = 0 and *s_e_* = 0.01. Here there is only one equilibrium point and it is stable. The (*s_e_*, *z*) subsystem is thus characterized by Class II excitability. As can be seen by comparing A to (B,C), the combined adaptation and inhibition has a strong effect on the amplitude of *s_e_* We note that if *g_ee_* is too small then the cubic nature of the *s_e_*-nullcline disappears and there is no excitability. We additionally note that with *g_ad_* ≈ 0, then when *g_ei_* falls below a critical value, there will be an additional stable fixed point with *s_e_* > 0.1 so that the local dynamics will not be excitable, but instead, it will be bistable.

**Fig 5.**
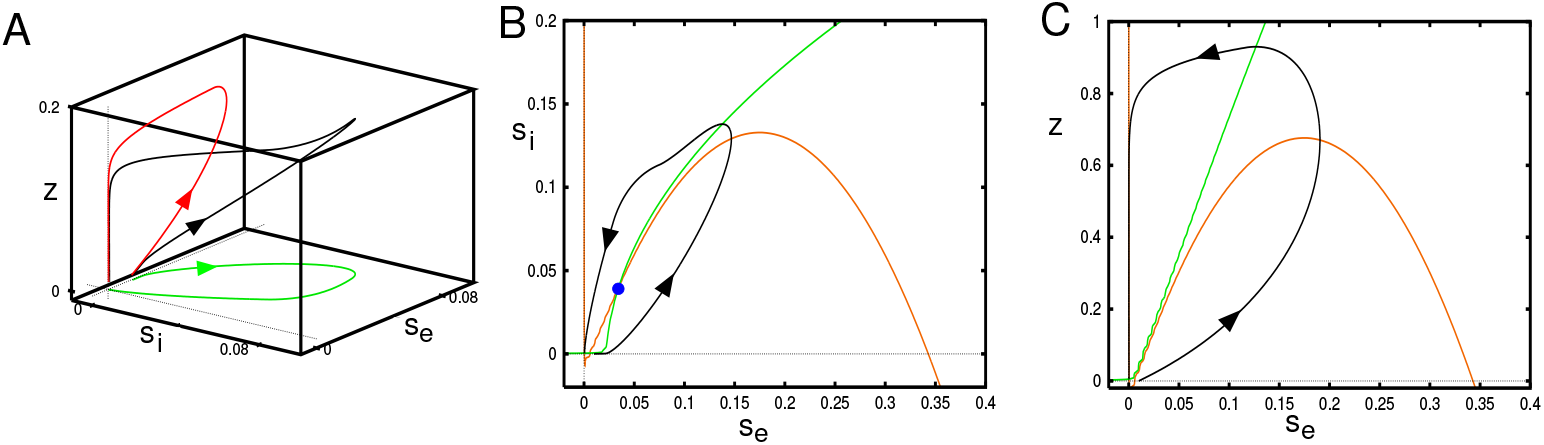
Excitability of the local equations (7). (A) Trajectory in (*s_e_*, *s_i_*, *z*) for *g_ei_* = 2, *g_ad_* = 0.25 with initial data (0.015, 0, 0). (B) (*s_e_, s_i_*)−plane when *g_ad_* = 0, *g_ei_* = 2 showing the excitatory (red) and inhibitory (green) nullclines (B) (*s_e_, z*) plane when *g_ad_* = 0.25, *g_ei_* = 0 showing the excitatory (red) and adaptation (green) nullclines

In sum, the mean field local kinetics show the hallmarks of an excitable system so that we expect that there can be robust travelling pulse waves in the spatially connected system.

### Smooth spatial system

We turn our attention to the full mean-field spatial model:

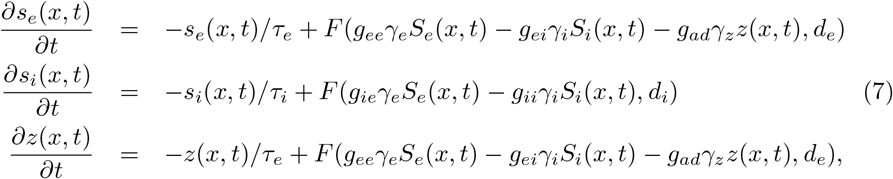

where

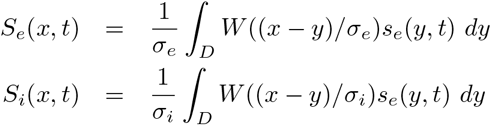

and *W* (*x*) is either t√he exponential, *W_E_*(*x*) = exp(−|*x*|)/2 or the Gaussian, 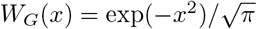 kernel. For purposes of analysis, *D*, the integration domain will be *R*, but for simulations, it will be a discretized finite domain. In what follows, we will fix all parameters except those that deal with the negative feedback, *g_ei_, g_ad_* and the spread of inhibition, *σ_i_* which has some very interesting effects on the stability of the waves.

To get a sense of the behavior of Eq. (7), we numerically solve the equations on a domain [0, 80] with a spatial discretization of dx = 0.2 and *σ_e_* = 1. We evoke a wave by setting *s_e_*(*x*, 0) = 0.2 for *x* ∈ [0, 4]. Fig. 6 shows the results of these simulations for the exponential (top row) and Gaussian (middle row) kernels. The figure also shows plots at *x* = 40 for *s_e_, s_i_, z* and *s_e_*(60, *t*) in the bottom row for the exponential kernel. The wave is faster for the exponential kernel since the spread of excitation decays slower than for the Gaussian. The left most column is the default set of parameters, *g_ei_* = 2, *g_ad_* = 0.25, *σ_i_* = 0.5, so that the spread of inhibition is half that of the excitation. With either of *g_ei_, g_ad_* removed, the excitation increases in magnitude and persists for a longer time. This can be seen clearer in the bottom plots. The velocity of the wave also increases as can be seen from the smaller slope. Increasing the spatial extent of inhibition to match that of the excitation (*σ_i_* = 1) has the effect of slowing the wave down. In addition, one can see a small oscillation near the onset of the wave in the exponential kernel case. We will spend some time exploring this oscillatory instability.

**Fig 6.**
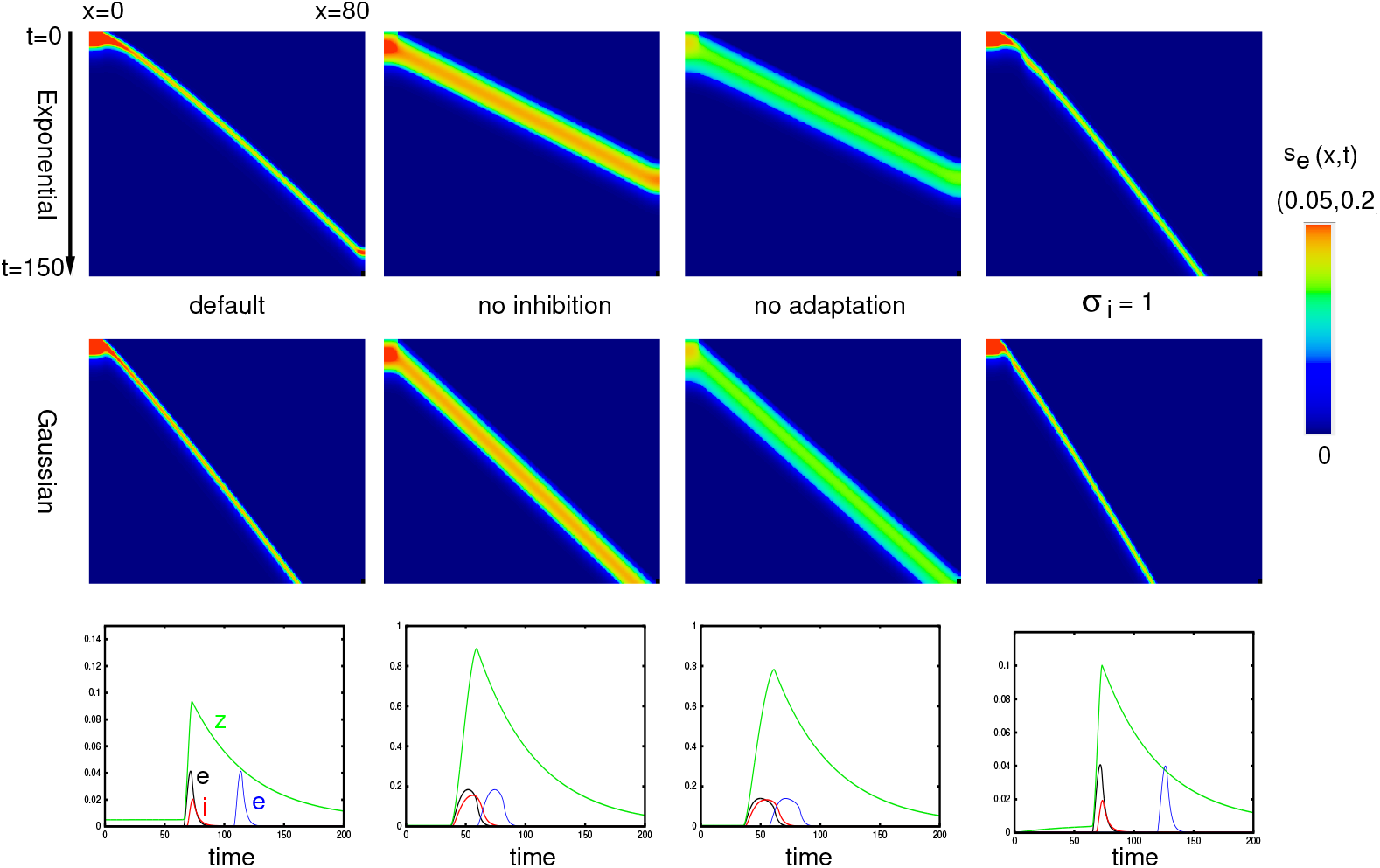
Plots of *s_e_*(*x, t*) for various parameters. Top row is the exponential kernel and second row is the Gaussian kernel. The default parameters are *g_ei_* = 2, *g_ad_* = 0.25, *σ_i_* = 0.5. The color scale has a maximum of 0.05 for the outer two plots and 0.2 for middle two plots. Bottom row shows *s_e_*(40, *t*), *s_i_*(40, *t*), *z*(40, *t*) and *s_e_*(60, *t*) of the corresponding parameters for the exponential kernel. Note the different vertical scales.

#### 0.1.1 Continuation and Reduction to a BVP

We would like to more systematically study the behavior of traveling wave solutions to Eq. (7) as we vary parameters. Thus, we convert to traveling wave coordinates, *ξ* = *x* + *ct*, so that Eq. (7) can be written as:

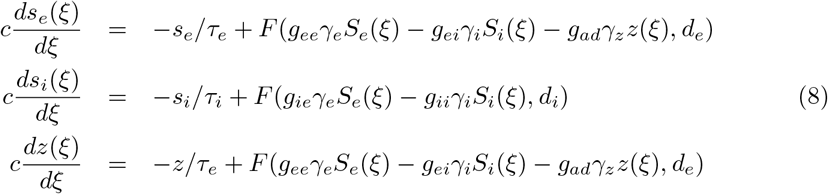

Let 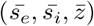 be the equilibrium value of the medium at rest. The solutions, (*s_e_*(*ξ*), *s_i_*(*ξ*), *z*(*ξ*)) must approach this equilibrium as *ξ* → ±∞. Unfortunately, this is an integro-differential equation and while there are some recent numerical tools for solving this type of equation [35], they are not yet well-developed and generally applicable. However, if we choose the exponential kernel, then we can readily invert the convolution to obtain a second order ODE and thus reduce Eq. (8) to an ODE. Specifically, if

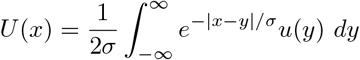

then

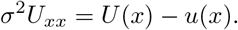

Thus we can write *S_e_*(*ξ*), *S_i_*(*ξ*) as a pair of ODEs and Eq. (8) becomes:

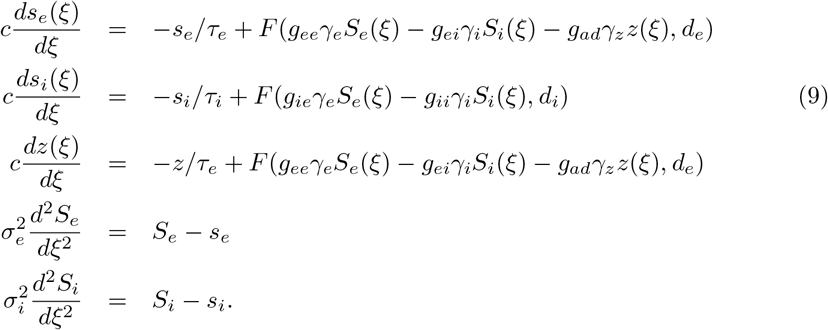

This is a 7-dimensional ODE for which we wish to find a homoclinic orbit that corresponds to the traveling pulse solution. Writing the *S_e_, S_i_* second order equations as a first order system, the equilibrium for 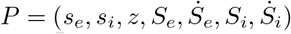 is 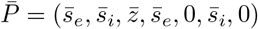. Linearizing around 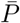, we find that there is a two-dimensional unstable manifold and a five-dimensional stable manifold. We use XPPAUT and AUTO to find the homoclinic orbit. More details on how we obtain a starting guess for AUTO are described in the methods.

We start by verifying that solutions to Eq. (9) are identical to the traveling waves obtained by simulation Eq. (7). Fig. 7 shows three examples with different values of (*g_ei_, g_ad_, σ_i_*). The plots essentially overlap so that we can be confident that the ODE approach provides the same results as integrating the full system of equations. Importantly, the ODE methods provide a tool that can determine the existence of these waves but it does not let us assess stability. We will explore the stability issue by solving Eq. (7) forward in time starting at a known wave solution.

**Fig 7.**
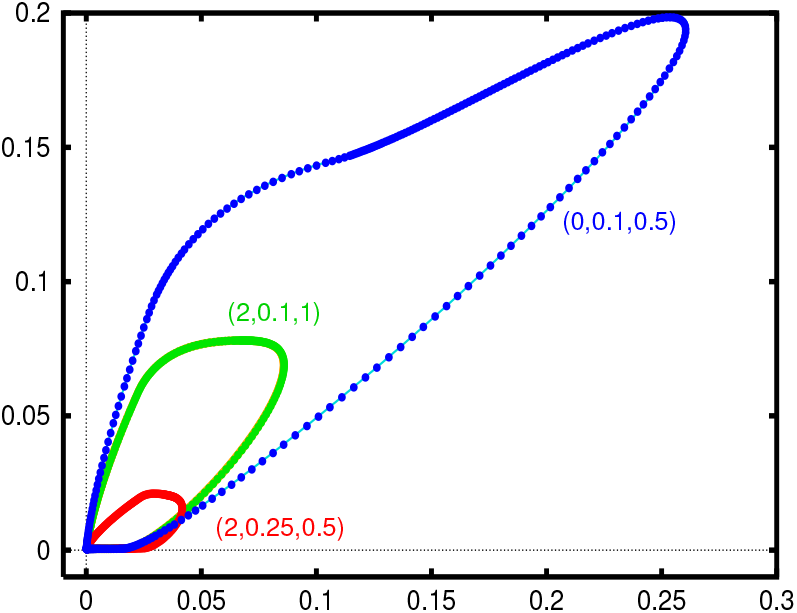
Comparison between solving Eq. (7) (thin lines) and the homoclinic computation of Eq. (9) (filled circles). Numbers in parentheses correspond to the values of (*g_ei_, g_ad_, σ_i_*).

#### 0.1.2 Dependence of the wave on parameters

In the next three figures, we hold two of the three parameters, (*g_ei_, g_ad_, σ_i_*) at fixed values and examine how the velocity varies as a function of the third parameter. We will also look at projections of solutions in the (*s_e_, s_i_*)− and (*s_e_, z*)− phaseplanes. Later when we study the piece-wise constant version of the model (with smooth nonlinearities replaced by step functions), we will also compute the width and velocity of the pulses and their stability.

Fig. 8A shows the behavior of the traveling pulse as *g_ei_* changes with *σ_i_* = 0.5 and three values of *g_ad_*. At the *g_ad_* = 0.25, the wave exists only if *g_ei_* is smaller than about *g_ei_* = *g_SN_* ≈ 5 where there appears to be a saddle-node bifurcation. For *g_ei_ < g_SN_*, there are two branches with a fast speed and a slow speed. Based on simulations of Eq. (7), we believe the fast waves are stable and the slow waves are unstable. (*s_e_, s_i_*) projections of solutions are shown panels (B,C). The amplitudes of both *s_e_, s_i_* increase as *g_ei_* decreases while the speed of the wave increases. For *g_ad_* = 0.1, we do not find the saddle-node point, at least for the range of *g_ei_* that we varied. For *g_ad_* = 0, the wave seems to stop at about *g_ei_* = 2.0. As we mentioned in section 2.2.1, at low values of *g_ad_, g_ei_*, the space-clamped system becomes bistable. We will explore this parameter region in the next section.

**Fig 8.**
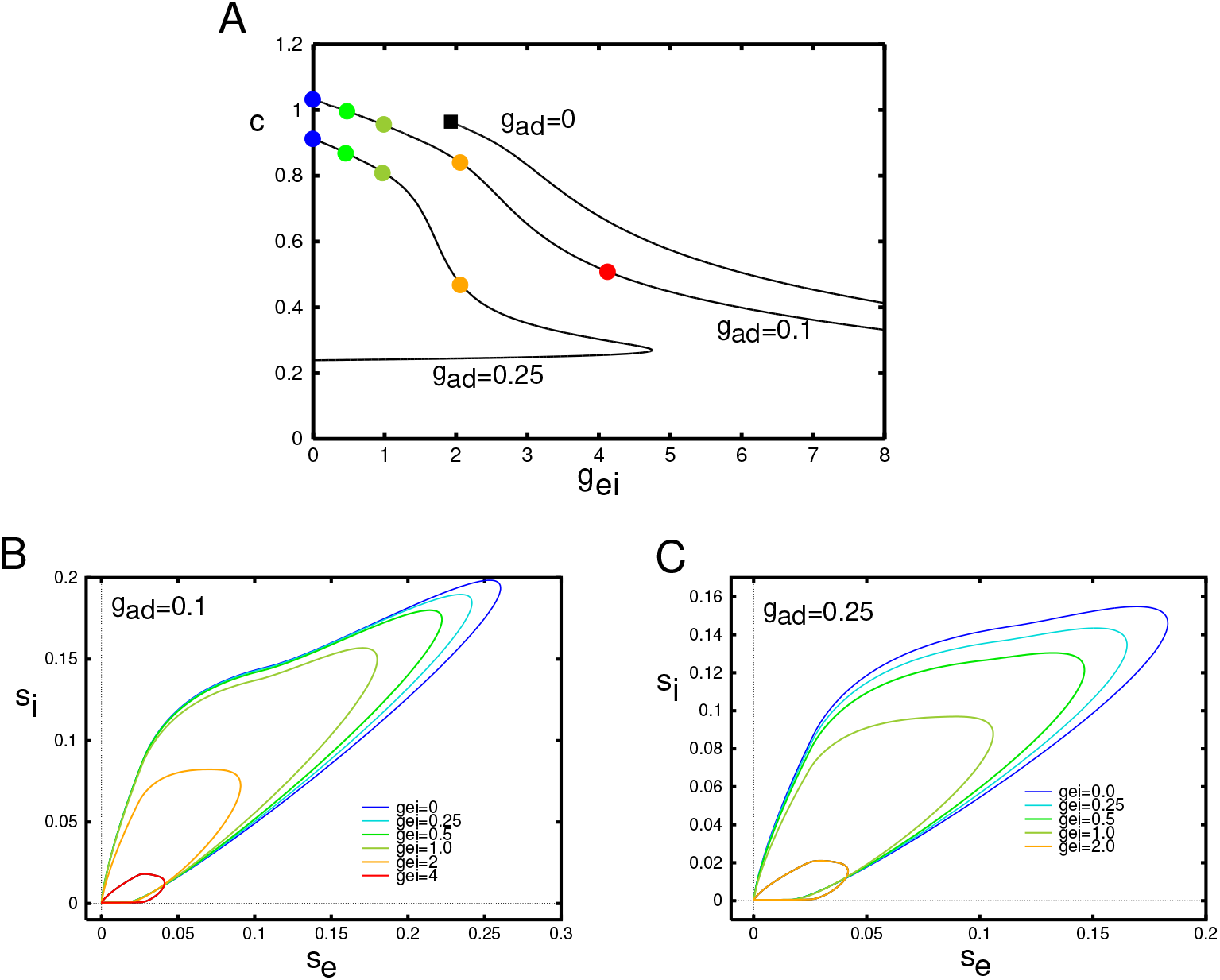
Behavior of Eq. (9) as *g_ei_* varies. (A) Velocity of the wave as a function of the strength of the inhibition onto the excitatory cells, *g_ei_* for three different levels of adaptation. Here *σ_i_* = 0.5 is fixed throughout. Colored circles correspond to the parameter values used in the projections onto the (*s_e_, s_i_*) phaseplanes in panels B,C. Small filled black square shows termination of the pulse to a front (see text).

Fig. 9 shows a similar plot where we fix *g_ei_, σ_i_* and vary *g_ad_*, the adaptation. The main difference is that adaptation has a much stronger effect, namely because of the fact that it decays much slower. A small increment in the firing rate is amplified by the adaptation. This can be seen by comparing the vertical axes in panels B,C; inhibition is considerably smaller than adaptation. Additionally, since *E_K_ < I_syn_*, the driving force of adaptation is also larger than that of inhibition. Because of the approximation we made for the adaptation, it is treated just like slow inhibition in the mean-field model. For all values of *g_ei_*, there are values of *g_ad_* such that there is a saddle node bifurcation and the wave ceases to exist. Based on the simulations of Eq. (7), we believe that the upper (“fast”) branch of solutions is the stable branch of solutions.

**Fig 9.**
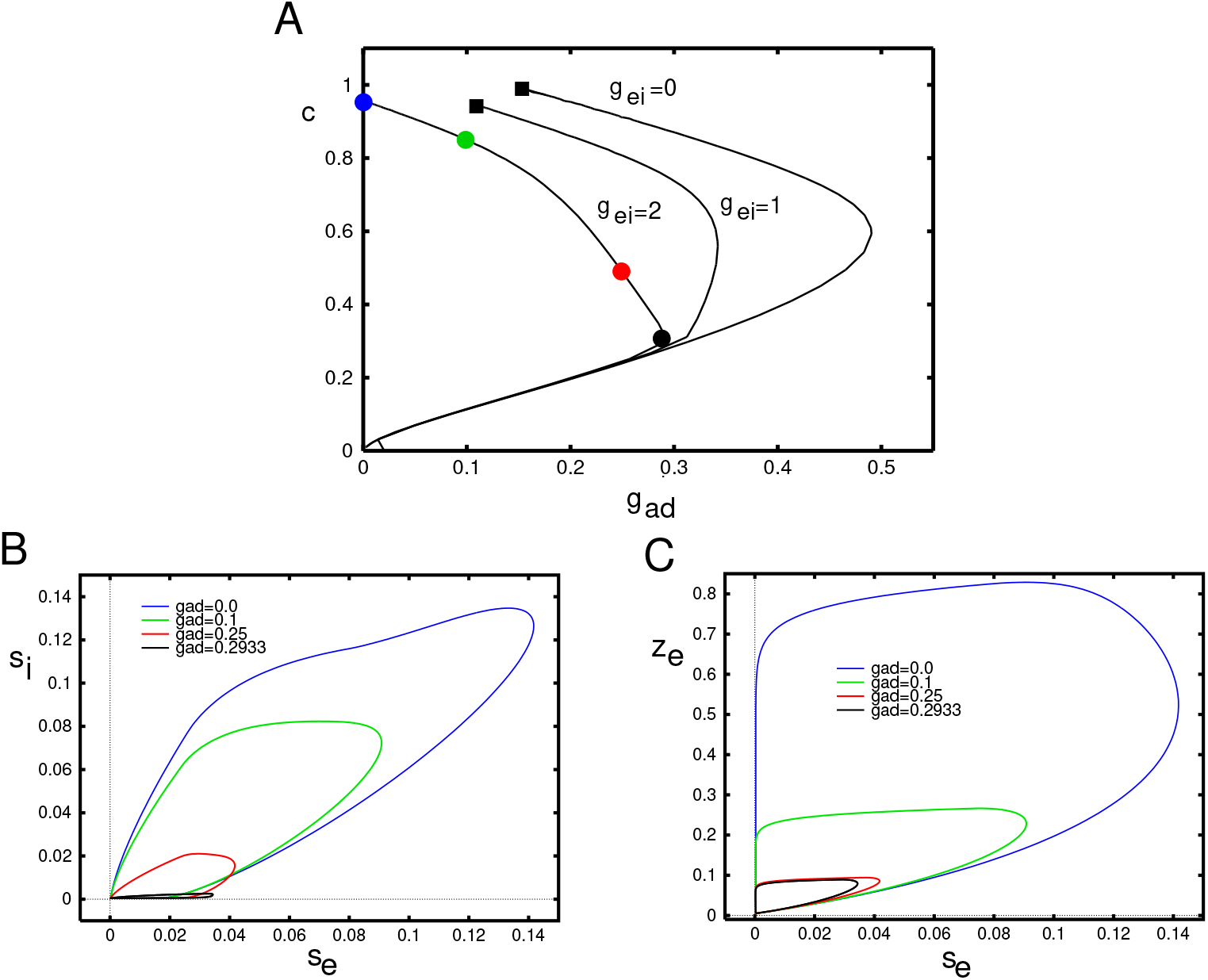
Behavior of Eq. (9) as *g_ad_* varies. (A) Velocity of the wave as a function of the adaptation, *g_ad_* for three different values of *g_ei_*. Here *σ_i_* = 0.5 is fixed throughout. Colored circles correspond to the parameter values used in the projections onto the (*s_e_, s_i_*)− and (*s_e_, z*)− phaseplanes in panels B,C. Small filled black squares as in Fig. 8.

In both Fig. 8 and 9, we see that when both *g_ei_* and *g_ad_* are small, the pulse ceases to exist (small filled black squares). Unlike the saddle-nodel bifurcations which mark the termination of any waves, here the pulse wave becomes a *front wave*. In the case of fronts, the local behavior (c.f. Fig. 5) is bistable. That is there is a stable low firing state and a stable high firing state. The wave connects the low state to the high state so that eventually the entire network is firing at a high rate. The transition from pulses to waves can be complex and has been well-studied in the reaction-diffusion literature [36].

Fig. 10 shows the wave behavior as a function of the spatial spread of inhibitions for three different pairs of (*g_ei_, g_ad_*). As would be intuitively expected, in all cases, the velocity decreases with the spread of inhibition. The wave appears to exist for *σ_i_* even three times greater than *σ_e_*. The shape of the wave in the (*s_e_, s_i_*)− phaseplane shows just a small effect of the inhibitory spread. However, we find that *σ_i_* induces some interesting effects on the stability of the traveling pulse. In the next part of the paper, we will simulate the waves via Eq. (7) to assess the stability of the solutions shown in this part of the paper. We find that for *g_ei_* large enough, the traveling wave loses stability as *σ_i_* increases, leading to an apparent Hopf bifurcation. The critical values of *σ_i_* are depicted by the filled black squares in panel A.

**Fig 10.**
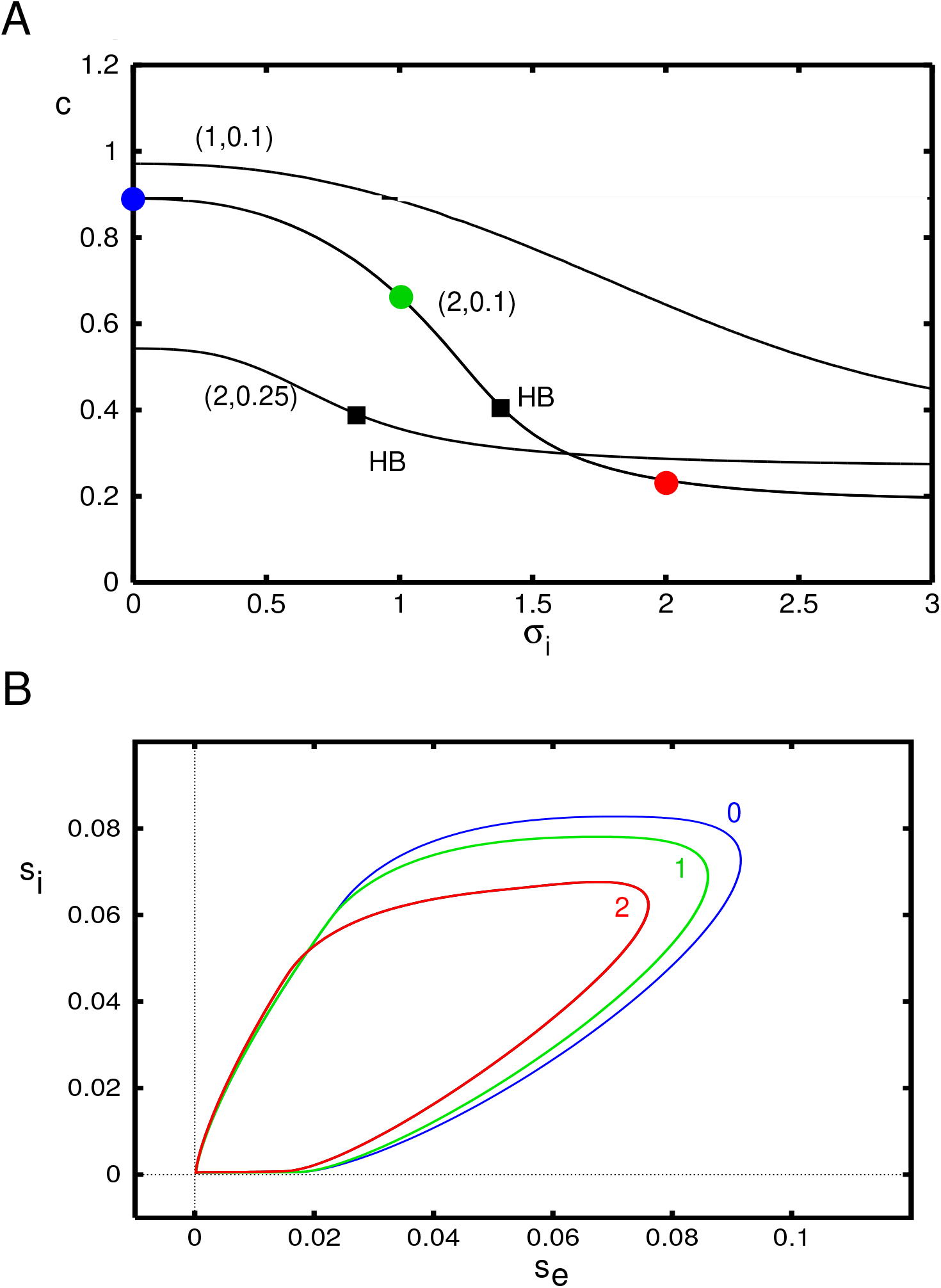
Behavior of Eq. (9) as *σ_i_* varies. (A) Velocity of the wave as a function of the spatial decay of inhibition, *σ_i_* for three different pairs (*g_ei_, g_ad_*). Filled circles correspond to the parameter values used in the projections onto the (*s_e_, s_i_*)− phaseplane in panel B. Squares mark apparent Hopf bifurcations of the waves.

#### 0.1.3 Instabilities of the wave

We have used the solutions obtained by our shooting methods as initial data in Eq. (7) and then solved the resulting equations forward in time to test stability. In the cases where *g_ei_* or *g_ad_* vary, but *σ_i_* = 0.5 is fixed, we have found that the solutions on the upper velocity curve appear to be stable. For example, when *g_ad_* = 0.1, we have increased *g_ei_* to 8 and find that the wave persists. However, this does not seem to be the case when we increase *σ_i_*, the spread of inhibition. Fig. 10 shows that the wave *exists* up to at least *σ_i_* = 3. On the other hand, fig. 11 (top) shows the result of starting near the exact traveling wave for three values of *σ_i_*. For *σ_i_* = 1.2 (recall that *σ_e_* = 1) one can start to see a bit of oscillation in the wave propagation, but it damps out and all eventually becomes the regular wave. When *σ_i_* = 1.48 the wave propagates but there is a spatiotemporal modulation of the wave. It appears that there is a Hopf bifurcation to an oscillatory modulated wave as the spread of inhibition increases. Destabilization of traveling pulses has also been reported in other work [19, 37] and appears to be through a similar mechanism. The modulated wave finally seems to break down once *σ_i_* gets too large (here, around *σ_i_* ≈ 1.54). This behavior is not confined to the exponential kernel, nor does it require a smooth firing rate function as can be seen in the middle and lower rows respectively.

**Fig 11.**
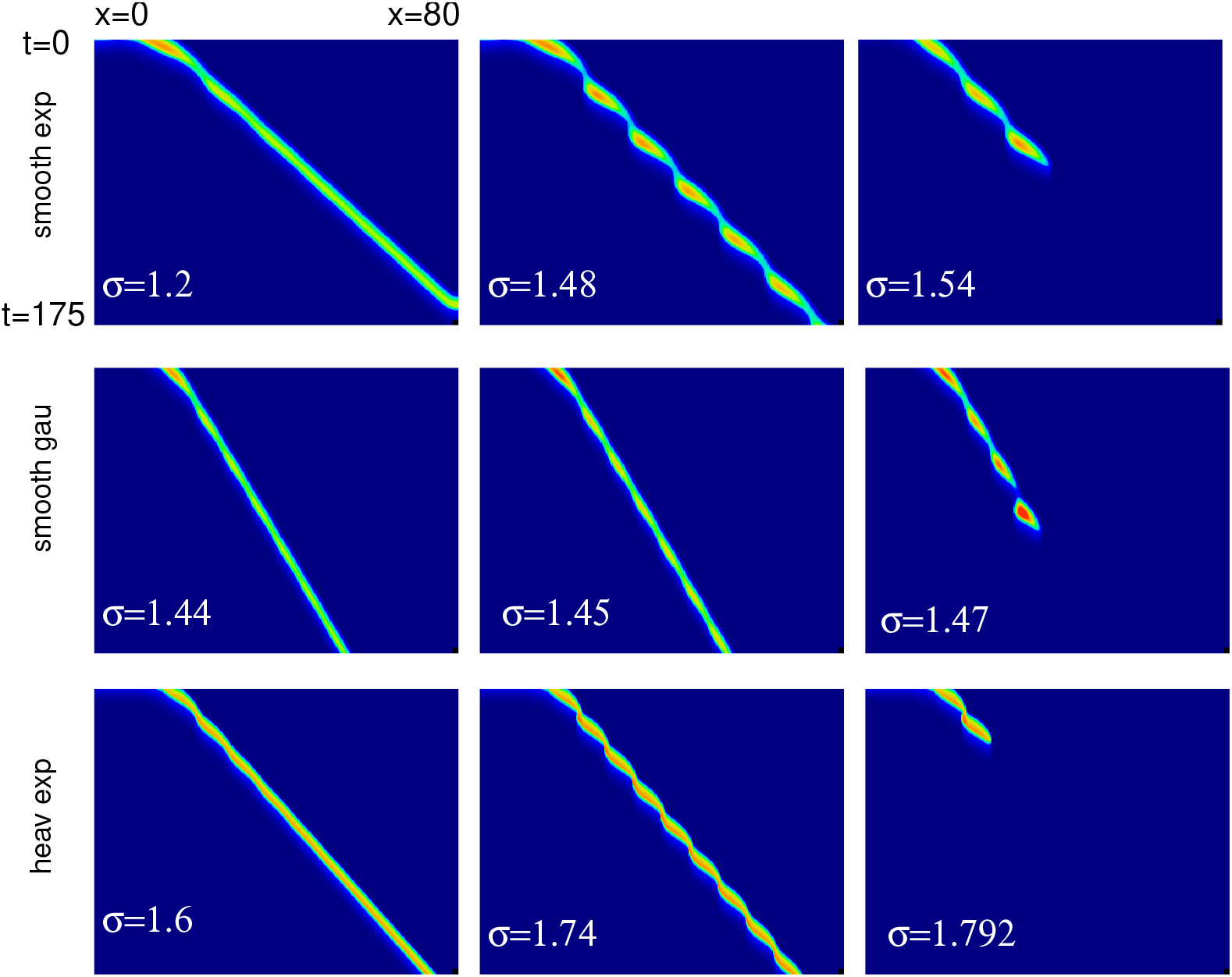
Simulations of the traveling pulse for Eq. (7) with (*g_ei_, g_ad_*) = (2.0, 0.1) for different values of *σ_i_*. Top row: smooth firing rate function (3) and exponential kernel; middle roe: smooth firing rate, Gaussian kernel; last row: Heaviside firing rate function, exponential kernel.

#### 0.1.4 Low adaptation

Additional bifurcations and pattern formation as well as multistability can be found in Eq. (7) when the adaptation is small or turned off. Fig. 12 shows some examples of the dynamics of the solitary traveling wave when *g_ad_* = 0 and *σ_i_* increases. (We remark that the same transitions are seen for *g_ad_* nonzero but sufficiently small.) In Panel A, the waves appears to “bounce” off the boundaries forming repeated zigzag waves. As *σ_i_* increases these waves develope periodic modulations (panel B) such that further increases in *σ_i_* break up into regular stripes (panel C). These stripes persist for larger *σ_i_*. They are large amplitude and, since the rest state remains asymptotically stable, they do not directly emerge as Turing patterns. These stripes coexist with the zigzag waves and the solitary traveling waves for a range of *σ_i_*. In panel D, we start at the striped initial conditions but lower *σ_i_* to 1.51 and observe the stripes break up into a series of traveling pulses.

**Fig 12.**
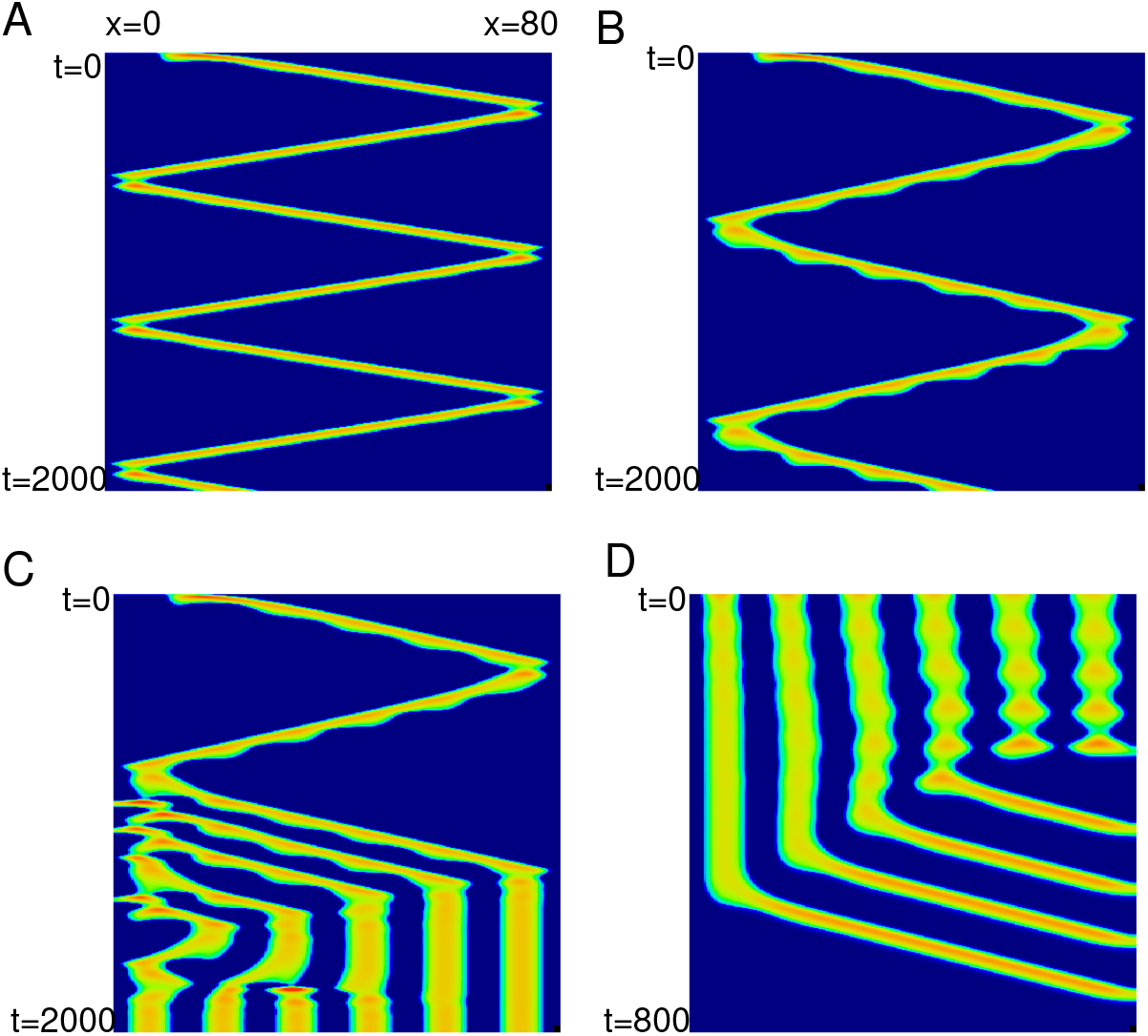
Complex dynamics without adaptation, *g_ad_* = 0. (A) *σ_i_* = 1.6; (B) *σ_i_* = 1.636; (C) *σ_i_* = 1.64; (D) *σ_i_* = 1.51.

#### 0.1.5 Heaviside approximation

In order to get some more insight into the behavior of Eq. (7), we will approximate the nonlinearity (3) by a step function and then rescale to put it into a simpler form that is suitable for analysis. The first step is to choose parameters, *a*_1_, *a*_2_, so that if we replace the smooth *F* (*G, d*) with *a*_1_*H*(*G* − *a*_2_), then, quantities such as the wavespeed and amplitudes are similar at some nominal parameter value. We pick (*g_ei_, g_ad_, σ_i_*) = (2, 0.1, 1), we choose *a*_1_ = 0.04, *a*_2_ = 0.12 so that the velocity of the waves matches that of the exponential kernel with smooth nonlinearity. Fig. 13 shows the comparison of the times series for *s_e_*(40, *t*) and phaseplane projections for the smooth and Heaviside function nonlinearities as and exponential and Gaussian kernels. Panels A,B show that while the heights do not match well (the step function saturates), the widths are roughly the same as is the velocity. For the Gaussion kernel, with our choice of parameters, the step function wave is much faster than the smooth wave. In panel B, we remove all the inhibition and increase *g_ad_* to 0.25. Here both the velocity and width still match and the results for the Gaussian kernel are more similar. In panels (C,D), the projections into the phaseplanes are shown. The plots are all qualitatively similar.

**Fig 13.**
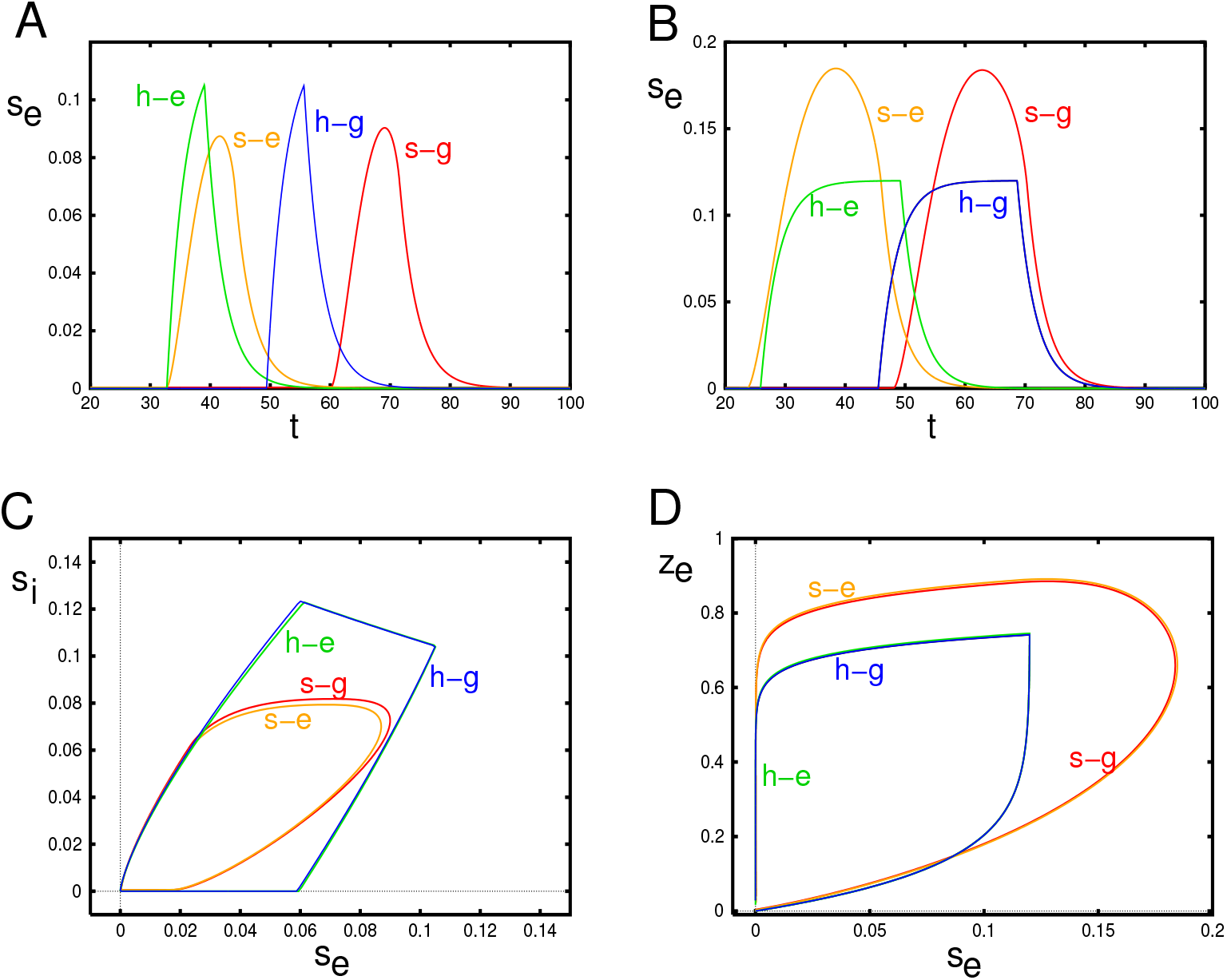
Comparison between the smooth model (s) and the Heaviside (h) models for the exponential (e) and Gaussian (g) kernels. In (A, C),(*g_ei_, g_ad_, σ_i_*) = (2, 0.1, 1); in (B,D) *g_ei_* = 0, *g_ad_* = 0.25 Time series plots show *s_e_*(40, *t*) and phaseplanes show *s_i_*(40, *t*) and *z*(40, *t*) on the *y*-axes.

#### 0.1.6 Analysis of the Heaviside model

As a first step, we replace *s_e_, s_i_, z* by 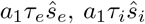, and 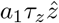 respectively and then drop the hats over the variables to obtain a “normalized” step function model:

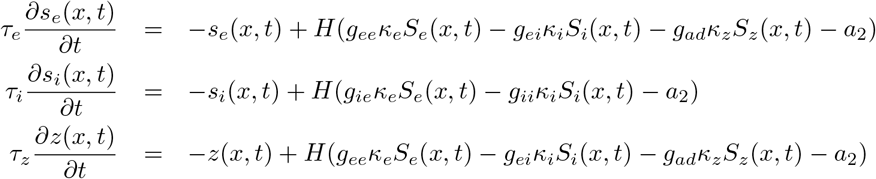

where *κ_β_* = *a*_1_*τ_β_γ_β_*. We replace *z*(*x, t*) in the adaptation by

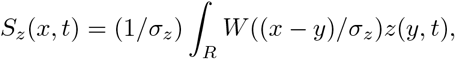

where we let *σ_z_* be very small. As we will see below, the reason for this is to avoid some technical difficulties when analyzing the stability. Now *s_e_, s_i_, z* all take values between 0 and 1. For our parameter choices *κ_e_* = 0.92, *κ_i_* = 0.37333, and *κ_z_* = 7.333333.

To study the existence and stability of traveling waves, we replace *x* by the moving variable, *ξ* = *ct* + *x*. Traveling waves are then just solutions independent of *t*. In this coordinate system, we obtain

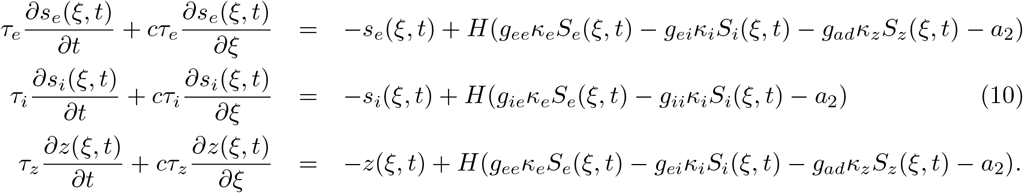

Steady states can be written as *s_e_*(*ξ, t*) = *u_e_*(*ξ*) etc where each *u_β_*(*ξ*) must vanish as *ξ* → ±∞. Let

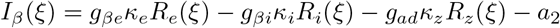

for *β* ∈ {*e*, *i*} and *R_e,i,z_*(*ξ*) are the convolutions of *u_e,i,z_*(*ξ*) with their respective kernels. (Note that there is no adaptation for the inhibitory cells.) The step function is 0 (1) if *I_β_*(*ξ*) is negative (resp., positive). Let *I_e_*(*ξ*) be positive for 0 < *ξ < a*, and *I_i_*(*ξ*) be positive for *b < ξ < d*. Our first goal is the solve for (*c, a, b, d*) whose values characterize the wave. (Note that since the waves are translation invariant, we can shift *ξ* so that *u_e_*(*ξ*) “turns on” at *ξ* = 0.) By continuity, we must have *I_e_*(0) = *I_e_*(*a*) = 0 and *I_i_*(*b*) = *I_i_*(*d*) = 0 along with the requirement that the derivatives of *I_e,i_*(*ξ*) with respect to *ξ* be nonzero at these crossings. Given (*c, a, b, d*) we can solve for *u_e,i,z_*(*ξ*) as they solve

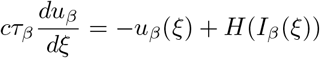

(where *I_z_*(*ξ*) = *I_e_*(*ξ*)). For example:

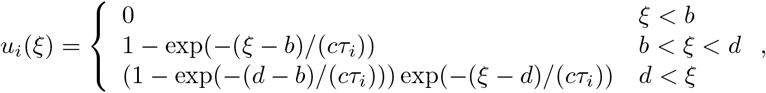

with similar expressions for *u_e_*(*ξ*) and *u_z_*(*ξ*). Given these functions, we can then evaluate the convolutions, *R_e,i,z_*(*ξ*) and thus obtain *I_e_*(*ξ*), *I_i_*(*ξ*) in terms of (*ξ, c, a, b, d*). Setting them to zero at the appropriate values of *ξ* gives us four equations in four unknowns:

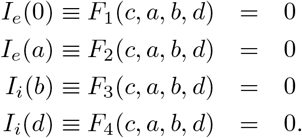

We get good guesses for (*c, a, b, d*) using the simulation of Eq. (7) (with the step function nonlinearity) and then using a root finder. The functions *F_j_* are complicated but easily obtained from the integrals using a symbolic algebra package.

Fig. 14 shows how various parameters related to the shape of the pulse vary as the three main parameters, *g_ei_, g_ad_, σ_i_* vary around their nominal values. In particular, the speed, *c* can be compared to the behavior of the smooth nonlinearity, Figs. 8–10. The speed decreases with any increases in these three parameters. Recall that *a* is the value of *ξ* where the input into the excitatory population falls back below threshold so that *a* is a surrogate for the width of the excitatory pulse. In fact, it is the distance from to time that excitation starts to the point that it reaches its peak. As intuitively expected, the width increases with reduced inhibition and adaptation. but has a nonmonotonic behavior with respect to *σ_i_*. For *ξ* (*b, d*), the input to the inhibition is above threshold; as with excitation, *b* marks the onset of firing for the inhibitory population and *d* marks its peak.

**Fig 14.**
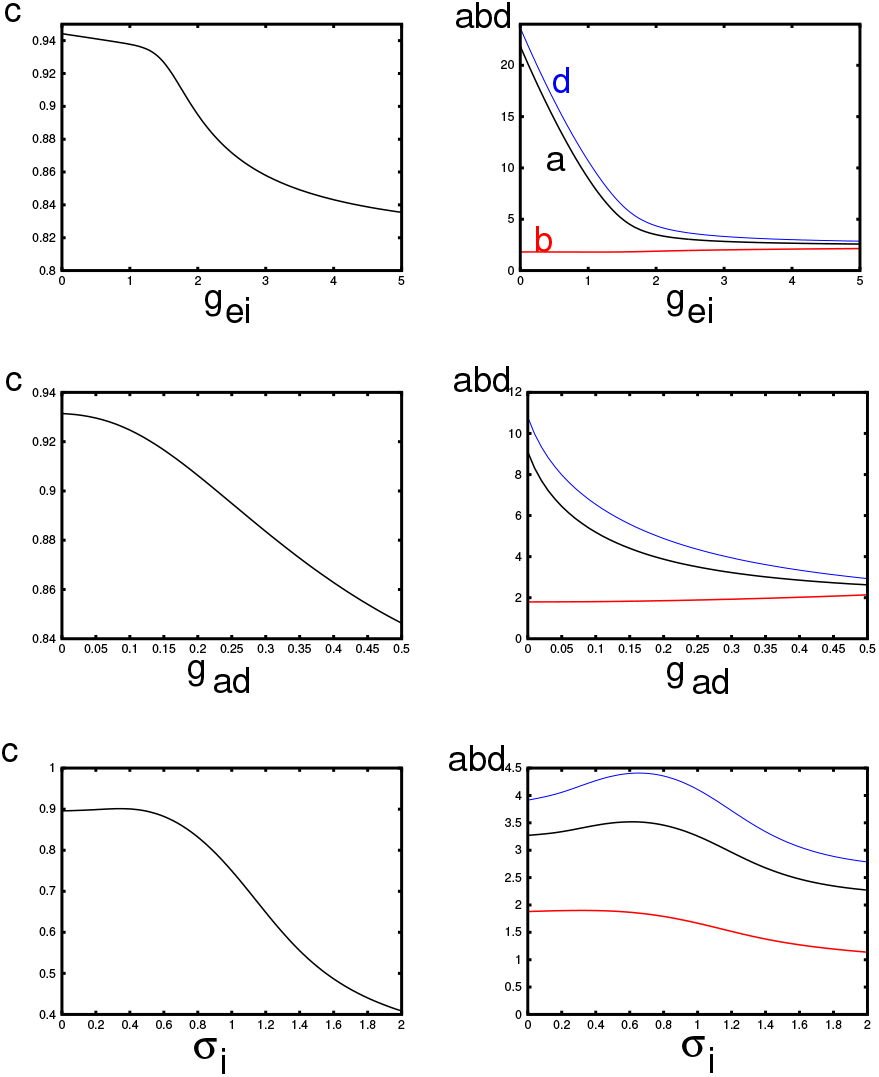
Properties of the traveling wave for the Heaviside firing rate function and the exponential kernel as parameters vary. The baseline values are (*g_ei_, g_ad_, σ_i_*) = (2, 0.25, 0.5). Here *c* is the wave speed, *a* is the width of the suprathreshold input to the excitatory population, *b* is the onset of the inhibition, and *d* is the offset. *b* − *d* is the width of the suprathreshold input to the inhibitory population.

#### 0.1.7 Stability of the wave

The big advantage of the Heaviside function approach is that it is possible to determine the stability of the waves. In order to do this, we need to formally linearize Eq. (10) about the traveling wave solutions, (*u_e_*(*ξ*), *u_i_*(*ξ*), *u_z_*(*ξ*)). Let *s_e_*(*ξ, t*) = *u_e_*(*ξ*) + exp(*λt*)*v_e_*(*ξ*) with similar expressions for *s_i_*(*ξ, t*), *z*(*ξ, t*) where *λ* and *v_β_*(*ξ*) are to be obtained. The wave will be unstable if ℜ*λ* > 0. Plugging this into Eq. (10) and retaining the linear terms, we obtain the following equations:

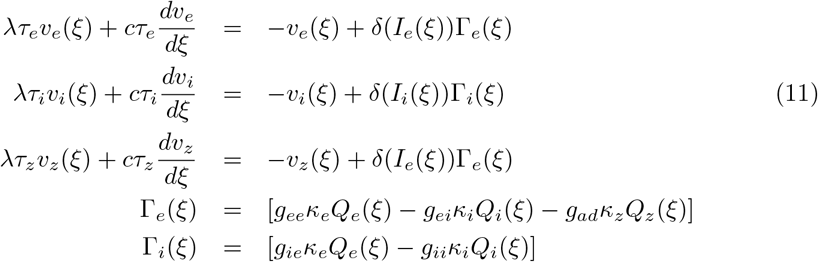

where *Q_β_*(*ξ*) are the convolutions of the variables, *v_β_*(*ξ*) with their respective kernels and the delta function is interpreted as:

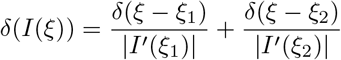

where *ξ*_1.2_ are the values of *ξ* where *I*(*ξ*) crosses 0. For excitation and adaptation, *ξ*_1_ = 0, *ξ*_2_ = *a* and for inhibition, *ξ*_1_ = *b, ξ*_2_ = *d*. Note that our “transversility” requirement implies that the derivatives of *I_β_*(*ξ*) at these points are nonzero. Before turning to the analysis of this equation, we first explain why we have convolved the adaptation with a sharp kernel. In each equation in the system (11) the delta functions are multiplied by Γ_*β*_(*ξ*). Consider the equation for *v_z_*(*ξ*). At *ξ* = 0 *v_z_*(*ξ*) must jump (since its derivative is a delta function). However, if we replaced *Q_z_*(*ξ*) with *v_z_*(*ξ*), then Γ_*e*_(*ξ*) would have a discontiuity at *ξ* = 0, so it is not clear how to evaluate the jump in *v_z_*(*ξ*) at *ξ* = 0. For this reason, we convolve *v_z_*(*ξ*) with a narrow kernel to assure that Γ_*e*_(*ξ*) is continuous at *ξ* = 0 (and *ξ* = *a*) and avoid this technical difficulty.

Consider *v_e_*(*ξ*) first. (The other two follow similarly.) We can write:

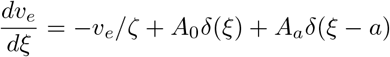

where *A*_0,*a*_ are the jumps for *v_e_*(*ξ*) and *ζ_e_* = *cτ_e_*/(*λτ_e_* + 1). The solution to this equation is:

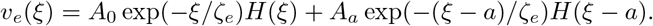

We similarly obtain:

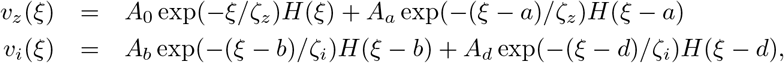

where *ζ_z,i_* are defined similarly to *ζ_e_*. The size of the jumps, *A*_0_,*_a,b,d_* is found by plugging 0, *a, b* of *d* into Γ_*β*_ (*ξ*) after dividing by the slopes 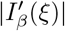 and *cτ_β_*. Let 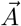 be the column vector (*A*_0_, *A_a_, A_b_, A_d_*)^T^. Then we obtain

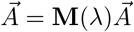

where **M**(*λ*) is a 4 × 4 matrix obtained from evaluating Γ_*β*_. For example, Γ_*e*_(0) depends on convolutions of *v_e,i,z_* evaluated at *ξ* = 0 and these depend linearly on *A*_0*,a,b,d*_. The coefficients multiplying the *A*_0_,*_a,b,d_* are then the entries of **M**(*λ*). We emphasize that each entry in **M** depends on the unknown eigenvalue, *λ*. The equation for 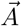 has a nontrivial solution if and only if

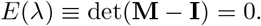

The complex function, *E*(*λ*) is called the Evans function. The zeros of this function are the eigenvalues for the linearized system. We set *σ_z_* = 0.001 in all the subsequent calculations; thus, *Q_z_*(*ξ*) is quite close to *v_z_*(*ξ*) but is continuous in *ξ*. We grid an area in the complex plane around *λ* = 0, and use MATLAB to compute *E*(*λ*). We then plot the zero contours of the real and imaginary parts of *E*(*λ*). Intersections of these contours are the eigenvalues. For each choice of parameters, we need to determine *c, a, b, d* and then compute the relevant integrals at the values of *ξ* ∈ {0, *a, b, d*}.

Recall Fig. 11 suggests that as *σ_i_* increases, the waves lose stability via a Hopf bifurcation. Thus, we consider the stability of the wave as we vary *σ_i_*. We fix *g_ei_* = 2, *g_ad_* = 0.25. Fig. 15 shows the results of this stability calculation. The top panel shows the contours of the real and imaginary parts of *E*(*λ*) as *λ* is varied in the region [−1, 0.2] + *i*[−1, 1]. The blue (red) curves show the real (imaginary) zero contours. They always intersect at *λ* = 0 corresponding to the translation invariance of the wave. For *σ_i_* = 1.5, there is a complex eigenvalue with a negative real part that is close to 0 (indicated by the filled circle). When *σ_i_* = 2.0, this intersection has moved over to the right half plane indicating that the wave is unstable. In Panel B, we follow this eigenvalue as we vary *σ_i_* and see that at *σ_i_* ≈ 1.8, the real part of the eigenvalue changes from negative to positive. At this point, the imaginary part is approximately, 0.45, indicating the possibility of a Hopf bifurcation. The numerical simulations we did above indicate that it is supercritical and a periodically modulated wave emerges for a narrow range of values of *σ_i_*. In Panels C and D, we find the critical value of *σ_i_* where R*λ* = 0 as we covary *σ_i_* and either *g_ei_* or *g_ad_*. In both cases, the dependence of *σ_i_* is nonmonotone. We were not able to make *g_ei_* (or *g_ad_*) too small before the calculation failed. In sum, our stability calculations indicate that as the spatial spread of inhibition increases, the traveling pulse loses stability through a complex eigenvalue. For a limited range of *σ_i_* beyond this critical value, there are periodically modulated waves. These waves eventually break up and the pulse can no longer propagate.

**Fig 15.**
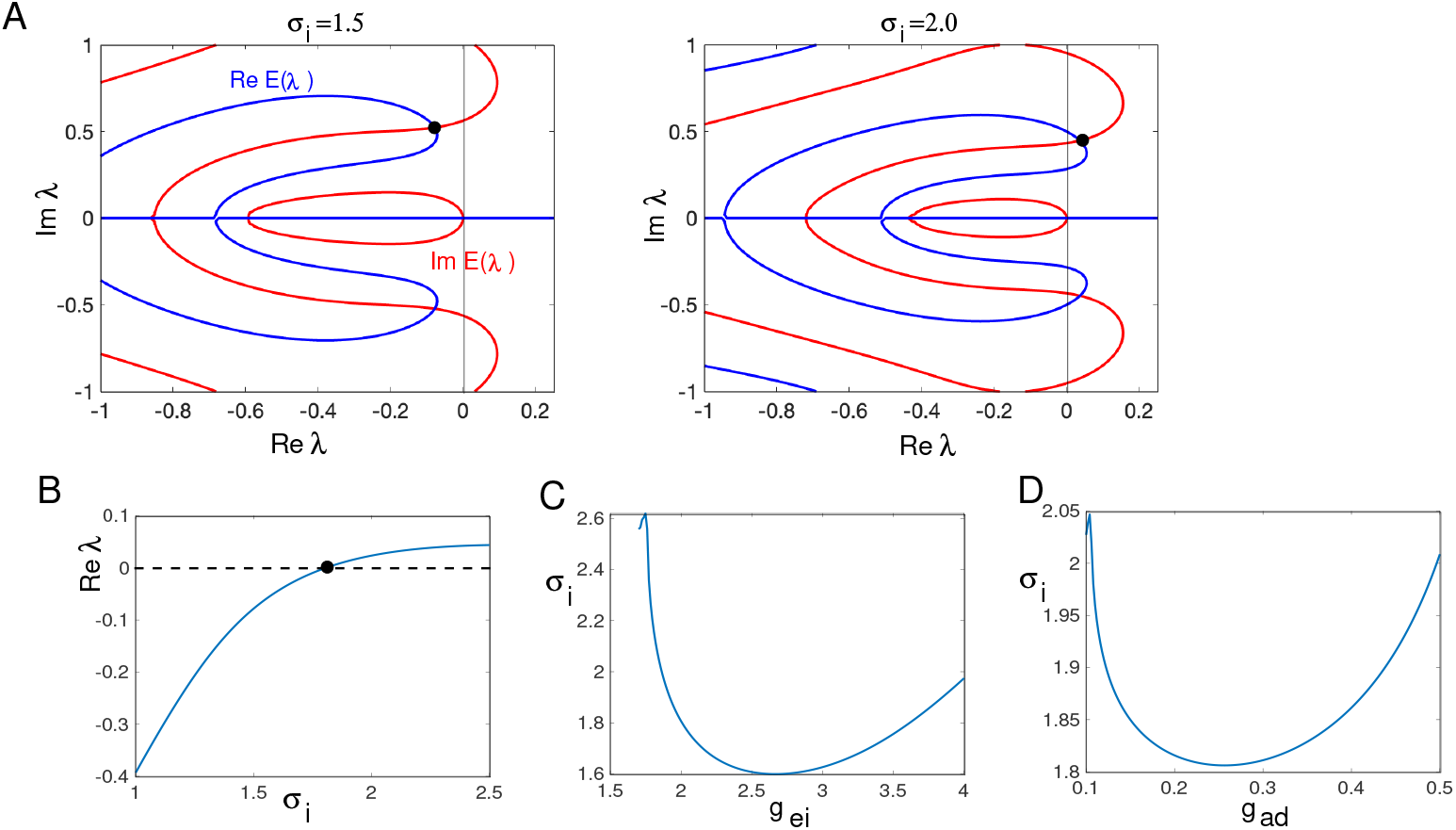
Stability of the traveling pulse. (A) The contours of the real and imaginary parts of the Evans function *E*(*λ*) whose intersections are the eigenvalues. *g_ei_* = 2.0, *g_ad_* = 0.25 and *σ_i_* is indicated in the figure. (B) Real part of the maximal nonzero eigenvalue as *σ_i_* varies. Zero crossing is a Hopf bifurcation. (C,D) Value of *σ_i_* where there is a Hopf bifurcation as *g_ei_* or *g_ad_* vary.

## Discussion

In this paper, we have explored the role of inhibition and adaptation on the control and propagation of traveling pulses in cortical networks. Starting with a spiking model, we showed that at high levels of recurrent inhibition, only about 10% of the excitatory neurons fire during a pulse and this is consistent with the small amplitude and somewhat irregular LFP seen in the experiment Fig. 1. Lowering the amount of inhibition led to both faster waves and much greater participation of the excitatory neurons. To gain better insight into how parameters change the properties of the waves, we derived a mean-field approximation for the spiking model. Using simulation and numerical shooting, we were able to study how the velocity and magnitude of the waves changed as parameters relating to negative feedback were changed. In particular, we found that without enough adaptation, as inhibition is reduced, the pulse wave is lost to a front. Conversely, as the strength or spread of inhibition increases, the pulse wave is lost through two mechanisms. First, it can be lost through a saddle-node bifurcation; as the inhibition increases, no wave exists. This loss of existence occurs at a finite value of velocity, *c* (c.f. Figs. 8,9). For this reason, we have not been able to get the same dynamic range of velocity as is seen in the experiments, (e.g. Fig. 1). The biggest dynamic range that we find is about 5-fold. The recent model by Gonzalez-Ramirez [12] has a similar range in velocities as we do. One possible explanation for the wide range in velocities seen in slice is that there are other sources of inhibition that are not directly dependent on the excitation. This “background inhibition” could act to increase the threshold to firing of the excitatory neurons which could lower the minimal velocity of the traveling waves. The second mechanism through which constant speed traveling waves are lost is through an apparent Hopf bifurcation. This instability leads to periodically modulated waves as seen in Fig. 11. In this case, we changed the “footprint” (spatial spread) of the inhibition, and eventually, the wave fails to propagate. In the Heaviside approximation, we show that changing either the strength of adaptation or inhibition can have a similar destabilizing effect, c.f. Fig. 15. Other interesting effects that we found in the smooth case are waves that reflect off the boundaries and transition to pattern formation (Fig. 12); the exact bifurcations which underly this are not completely resolved.

Our present work is similar to the paper by Gonzalez-Ramirez et al [12] in that both models study waves in networks with excitatory and inhibitory neurons in addition to a slow adaptation that serves to suppress run-away excitation. Their adaptation is linear whereas ours is nonlinear; both forms of adaptation have an large effect at damping the strong excitation. One main difference between our work and theirs is in the effects of inhibition. In their paper, the main source of control of activity is the adaptation while in ours it is the inhibition. Because adaptation is slow, it cannot control the magnitude of the excitation so that the waves that occur in adaptation-dominated networks are much broader and involve almost all of the excitatory neurons. This can be seen in or spiking simulations in Fig. 2 (*g_ei_* = 0) and in Fig. 3B. In our work the inhibition has a considerable effect on controlling the width of the wave in contrast to [12] (c.f. Fig. 7 in their paper). In their model, inhibition increased the wave speed while in ours and in the experimental results in Fig. 1, velocity decreased. They found that the inhibition had to be 10 fold slower than the excitation in order to attain propagation; in our model the time scales of excitation and inhibition were similar. One explanation for these different results is that they were primarily interested in seizure-type waves while our focus as been on sensory evoked waves. The underlying assumptions of the two different models are likely to be different. Their approach to varying parameters is also quite different from ours as throughout the paper they seek analytic expressions for the wave. As a consequence, they treat width and velocity as parameters in order to obtain parametric expressions for other parameters such as the strength of inhibition. We numerically compute velocity and width directly by continuation so we can compare the wave shape and velocity to the parameters relevant to inhibition. Finally, we found a number of bifurcations to more complex wave forms (e.g. periodically modulated waves) as we varied the spatial spread and strength of inhibition which provided us with information on how the waves cease to propagate.

In earlier experimental work [9, 38], these authors show that the ability of waves to initiate and propagate depends on the feedback between layers 2/3 and 5 in the cortex. The dense recurrent excitation in layer 5 coupled with the longer range excitatory interactions underly the initiation and propagation of traveling waves in normal and disinhibited cortex. Thus, a natural extension of the present work is to create a two-layer network and explore how strong recurrent connections in one layer (5) interact with weaker but longer-range connections in the other (2/3) to produce propagation.

In conclusion, in this paper we have tried to explore how inhibitory feedback and intrinsic adaptation work together to control the propagation of waves through space. Sensory evoked waves are one possible mechanism for transmitting information in one cortical area or region to another. Recurrent excitation is required to maintain the transmission in a robust manner. However, this strong excitation must be controlled. Our analysis and simulations suggest ways in which inhibition can keep the activity at reasonable levels while at the same time allowing for long distance propagation.

